# Enzymatic and structural characterization of HAD5, an essential phosphomannomutase of malaria parasites

**DOI:** 10.1101/2021.07.03.451008

**Authors:** Philip M Frasse, Justin J Miller, Ebrahim Soleimani, Jian-She Zhu, David L Jakeman, Joseph M Jez, Daniel E Goldberg, Audrey R Odom John

**Affiliations:** Division of Infectious Diseases, Departments of Medicine and Molecular Microbiology, Washington University School of Medicine, St. Louis, Missouri, USA; Department of Biology, Washington University in St. Louis, St. Louis, MO 63130, USA; College of Pharmacy, Dalhousie University, Halifax, Nova Scotia, Canada; Department of Chemistry, Razi University, Kermanshah, Iran; Department of Chemistry, Dalhousie University, Halifax, Nova Scotia, Canada; Division of Infectious Diseases, Department of Pediatrics, Children’s Hospital of Philadelphia, University of Pennsylvania, Philadelphia, Pennsylvania, USA

**Keywords:** Malaria, parasite, glycosylphosphatidylinositol (GPI) anchor, carbohydrate metabolism, drug development, haloacid dehalogenase (HAD), phosphomannomutase, crystal structure

## Abstract

The malaria parasite *Plasmodium falciparum* is responsible for over 200 million infections and 400,000 deaths per year. At multiple stages during its complex life cycle, *P. falciparum* expresses several essential proteins tethered to its surface by glycosylphosphatidylinositol (GPI) anchors, which are critical for biological processes such as parasite egress and reinvasion of host red blood cells. Targeting this pathway therapeutically has the potential to broadly impact parasite development across several life stages. Here, we characterize an upstream component of GPI anchor biosynthesis, the putative phosphomannomutase (EC 5.4.2.8) of the parasites, HAD5 (PF3D7_1017400). We confirm the phosphomannomutase and phosphoglucomutase activity of purified recombinant HAD5. By regulating expression of HAD5 in transgenic parasites, we demonstrate that HAD5 is required for malaria parasite egress and erythrocyte reinvasion. Finally, we determine the three-dimensional crystal structure of HAD5 and identify a substrate analog that specifically inhibits HAD5, compared to orthologous human phosphomannomutases. These findings demonstrate that the GPI anchor biosynthesis pathway is exceptionally sensitive to inhibition, and that HAD5 has potential as a multi-stage antimalarial target.

## INTRODUCTION

Malaria remains an enormous global health burden for much of the world, resulting in over 200 million infections and 400,000 deaths every year, the majority of which are in children under the age of five(1). One of the primary barriers to effective malaria treatment and control is the emergence of resistance to all approved antimalarial chemotherapeutics(2, 3), prompting an urgent call for the development of new therapies and the identification of novel drug targets. Malaria is caused by apicomplexan parasites of the genus *Plasmodium,* primarily the species *Plasmodium falciparum*. *P. falciparum* has a complex life cycle, in which parasites develop in a mosquito host, are deposited into a human during a mosquito blood meal, and migrate to the liver where they infect hepatocytes. Parasites are released from the liver into the bloodstream and begin an asexual cycle of replication within red blood cells, occasionally branching off into sexual stage gametocytes that can be taken up by mosquitoes to start the cycle anew(4).

This complex, multi-host life cycle has stymied efforts to develop therapeutics and vaccines to eradicate this disease, as it has been difficult to identify effective vaccine and therapeutic targets that span multiple life stages. In recent years, the push for new therapeutics and vaccines has focused on a strategy of developing transmission-blocking vaccines and therapies, which impair the development or viability of gametocytes or target the mosquito vector itself, thus preventing vector-borne transmission(5–7). Our goal is therefore to identify novel targets for antimalarial therapeutics that are not only essential for intraerythrocytic growth of the parasite, but are also essential for sexual- and/or mosquito-stage parasites, indicating their potential in transmission blocking strategies.

Malaria parasites are highly metabolically active(8, 9), and several known antimalarials target unique and essential metabolic processes in the parasites(10–14). Metabolic enzymes have great potential for chemical inhibition, as substrate analogs can be rationally designed and developed as potential inhibitors, making these enzymes well-suited as “druggable” targets(15). We therefore sought a metabolic enzyme that is expressed and essential during multiple life stages of *Plasmodium falciparum*. These constraints narrowed our search to the upstream steps of glycosyl phosphatidylinositol (GPI) anchor synthesis.

GPI anchors are an essential component of all life stages of *Plasmodium falciparum*. In intraerythrocytic parasites, these glycolipid anchors tether several essential proteins to the parasite plasma membrane prior to red blood cell egress and reinvasion(16). Most abundant amongst these GPI-anchored proteins (GPI-APs) is merozoite surface protein 1 (MSP1)(16–18). Proper MSP1 localization and processing is necessary for parasites to egress from the erythrocyte, and mature MSP1 anchored to the surface of free merozoites facilitates the binding and invasion of new red blood cells. In the absence of the GPI-anchoring C-terminus of MSP1, parasites are defective in their ability to egress(19), and antibodies developed against MSP1 prevent merozoite reinvasion(20). Several other GPI-APs are also involved in parasite egress and invasion, including other merozoite surface proteins (MSPs) and rhoptry-associated membrane antigen (RAMA)(16, 18, 21). GPI anchored proteins are also expressed in other life stages of *P. falciparum*. Gamete- and ookinete-stage GPI anchored proteins include Pfs25 and Pfs230, which are considered as possible vaccine candidates(22–25), while circumsporozoite protein (CSP), the critical antigen of the RTS,S and R21 vaccines, is an essential GPI-anchored protein of the sporozoite stages(26–30). Thus, it is clear that successful targeting of GPI anchor biosynthesis would not only effectively treat symptomatic blood-stage infection, but may also block transmission of the parasites at multiple stages.

One enzyme of the GPI anchor biosynthesis pathway that has yet to be characterized in *P. falciparum* is the putative phosphomannomutase, HAD5 (PMM; PF3D7_1017400). Phosphomannomutases (PMMs) are responsible for the conversion of mannose 6-phosphate (M6P) to mannose 1-phosphate (M1P), the precursor to GDP-mannose. GDP-mannose is then converted to dolichol-p-mannose, which is the building block for incorporating mannose into glycolipids. In asexual *P. falciparum*, the dominant mannosylated glycolipids are GPI anchors(31), as *N*-glycans in these parasites only contain *N*-acetyl glucosamine(32–34), so targeting of mannose metabolism is predicted to specifically inhibit GPI anchor synthesis. HAD5 is also predicted to be essential(35), and transcriptomic studies show its expression during the blood stage(36, 37) and sexual stages(38), making it a potential multi-stage antimalarial drug target. In this study, we characterize the putative phosphomannomutase of *P. falciparum*, HAD5, demonstrating its essentiality for parasite growth and its potential for specific targeting by future antimalarial therapies.

## RESULTS

### HAD5 is a phosphomannomutase

HAD5 (PF3D7_1017400) has been annotated as a putative phosphomannomutase through homology to other known phosphomannomutases. Phosphomannomutases are responsible for interconverting M6P and M1P to generate M1P for downstream glycolipid production, most notably GPI anchors in *P. falciparum* (Fig. 1A)(39–41). To determine the biochemical function of HAD5, we purified recombinant HAD5 (Fig. 1B) and examined its hexose phosphate mutase activities (Fig. 1C, S1). We found that HAD5 was active in the phosphomannomutase assay (0.56 ± 0.48 µmol/min/mg), with even more robust activity (7.67 ± 0.06 µmol/min/mg) upon addition of the known co-factor glucose-1,6-bisphosphate (G-1,6-P)(41), which is comparable to the specific activity of other phosphomannomutases(42, 43). Furthermore, as has been seen for other phosphomannomutases(43, 44), HAD5 exhibits some promiscuity in its substrate preference, as it also displays phosphoglucomutase activity at 0.31 ± 0.03 µmol/min/mg. Comparing the catalytic efficiencies of HAD5 toward phosphomannose and phosphoglucose suggests that phosphomannomutase activity is the dominant enzymatic function of HAD5, with a roughly 4-fold higher catalytic efficiency (k_cat_/K_m_) compared to its phosphoglucomutase activity (Table 1).

**Figure 1.**
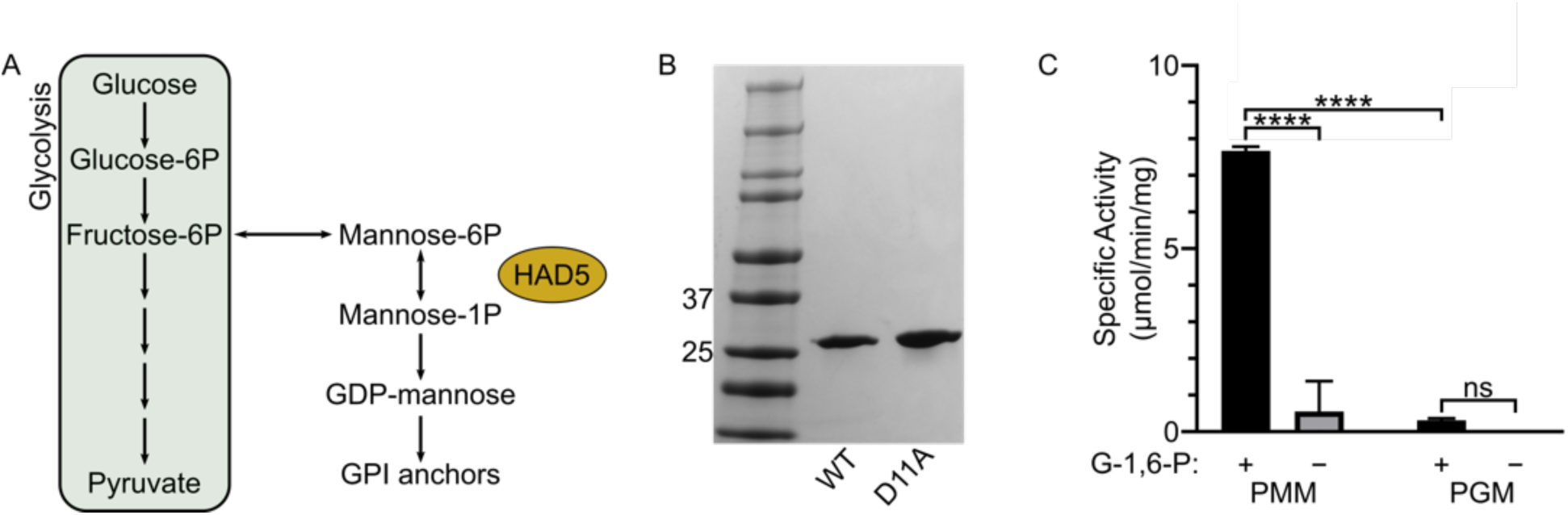
HAD5 is a bifunctional phosphomannomutase/phosphoglucomutase. **(A)** Schematic of phosphomannomutase’s role in metabolism. Phosphomannomutases like HAD5 interconvert mannose 6-phosphate and mannose 1-phosphate, providing the latter for downstream glycolipid production and synthesis of GPI anchors in *P. falciparum*. **(B)** SDS-PAGE gel of recombinant wild-type (WT) HAD5 and a catalytically inactive mutant (D11A). **(C)** Displayed are the mean ± standard error of the mean (SEM) of HAD5 activity across three independent trials, with D11A activity subtracted as background. Abbreviations: G-1,6-P, glucose-1,6-bisphosphate; PMM, phosphomannomutase assay; PGM, phosphoglucomutase assay. *p*-values were determined using an ordinary two-way ANOVA (Tukey’s test for multiple comparisons, α= 0.05). *\*\*\*\*p<*0.0001; ns = not significant.

**Table 1.** Phosphomannomutase and phosphoglucomutase activity of HAD5. Displayed are the mean ± SEM of three independent trials for the kinetic parameters of HAD5 converting M6P to M1P (PMM) or G1P to G6P (PGM).

| Enzyme Assay | $k_{cat}$ (s<sup>-1</sup>) | $K_m$ (μM) | $k_{cat}/K_m$ (M<sup>-1</sup>s<sup>-1</sup>) |
| --- | --- | --- | --- |
| **PMM** | 1.7 x 10<sup>-4</sup><br>± 1.9 x 10<sup>-5</sup> | 32 ± 3.6 | 5.4 |
| **PGM** | 7.8 x 10<sup>-6</sup><br>± 1.2 x 10<sup>-6</sup> | 5.2 ± 1.4 | 1.5 |

### HAD5 is essential for intraerythrocytic parasite growth

To assess the essentiality of HAD5 in intraerythrocytic parasite stages, we used a previously described conditional knockdown system in cultured asexual *P. falciparum*(45, 46) (Fig. 2A). We placed a Tet repressor-binding aptamer array at the 3’- and 5’-end of the endogenous HAD5 locus by CRISPR/Cas9-mediated integration. In parasite integrants, the presence of anhydrotetracycline (aTc) promotes HAD5 translation, while washing out aTc leads to inhibition of translation, and we term this conditional knockdown strain “HAD5^KD^”. Immunoblotting confirms substantial reduction in cellular abundance of HAD5 in the absence of aTc (Fig. 2B). In HAD5^KD^ parasites, 0 nM aTc conditions led to an absence of growth, whereas addition of aTc promoted growth in a dose-dependent manner (Fig. 2C), indicating that HAD5 is essential for asexual growth of *P. falciparum*.

**Figure 2.**
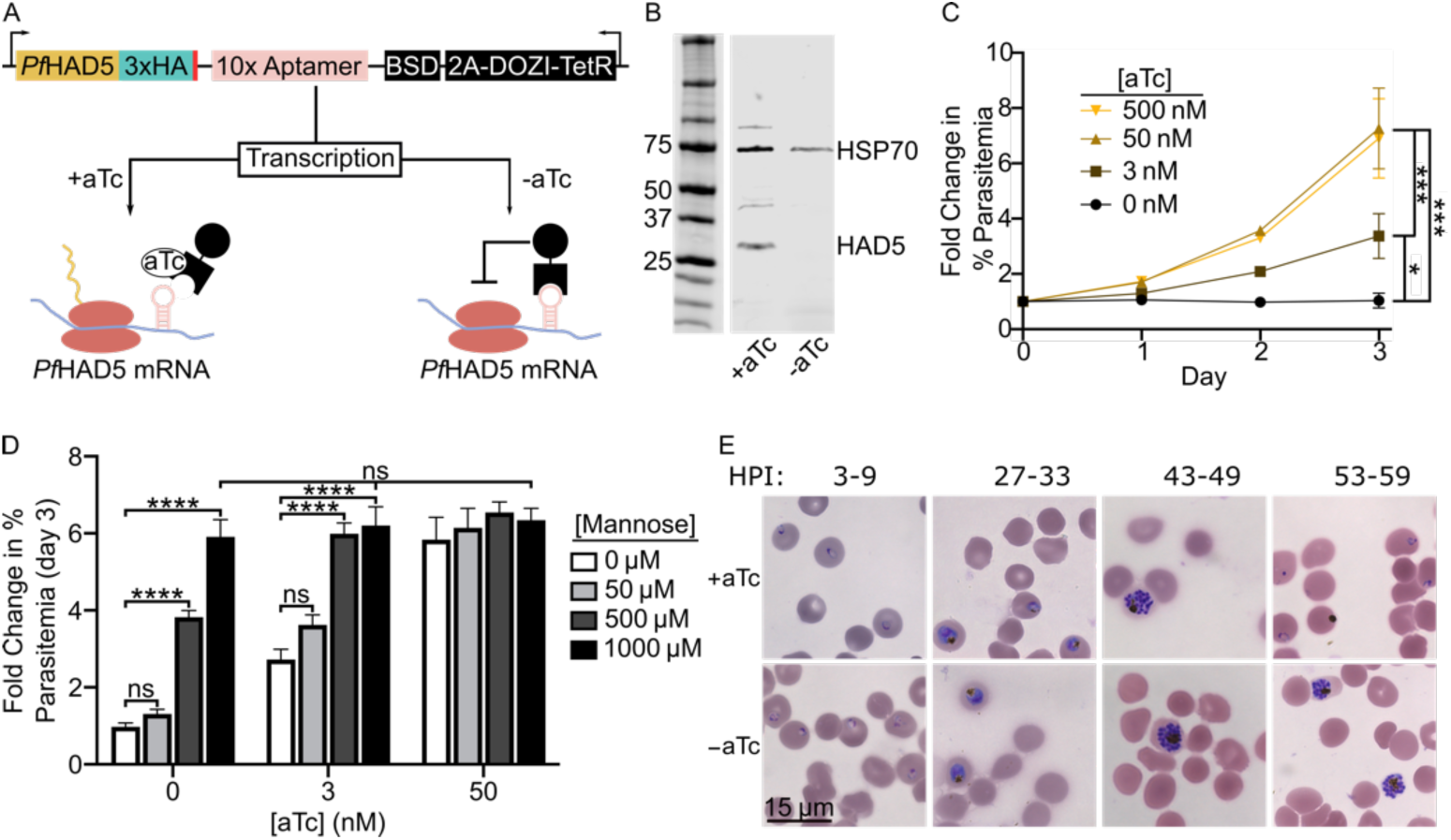
HAD5 is essential for intraerythrocytic parasite growth. **(A)** Schematic of the regulatable knockdown system introduced at the native locus of *Pf*HAD5(45, 46). **(B)** Western blot of transgenic HAD5^KD^ parasite lysate ±aTc, using α-HA to detect HAD5 and α-HSP70 as a loading control. Removal of aTc results in successful knockdown of HAD5. Approximate expected protein masses: *Pf*HSP70, 74 kDa; *Pf*HAD5-3xHA, 32 kDa. **(C)** Fold-change in parasitemia over time of HAD5^KD^ parasites grown in varying concentrations of aTc. Data represent mean ± SEM of three independent experiments with technical duplicates. Significance was determined by one-way ANOVA with Fisher’s LSD; \**p*=0.03; \*\*\**p*<0.001. **(D)** Fold change in parasitemia of HAD5^KD^ parasites grown in varying aTc concentrations with D-mannose rescue. Significance was determined by ordinary two-way ANOVA with Tukey’s correction for multiple comparisons. *\*\*\*\*p<*0.0001, ns = not significant. Data represent mean ± SEM of three independent experiments with technical duplicates. **(E)** Bright-field images of Giemsa-stained thin-smear synchronized parasites over time. HPI = hours post-invasion.

Because phosphosugar mutases often utilize more than one substrate, we examined whether the phosphomannomutase activity of HAD5 was responsible for its essential function in asexual parasites. Attempts to chemically rescue parasite growth with hexose phosphates such as M6P and M1P were unsuccessful (Fig. S2A), as was expected due to the impermeability of the erythrocyte and parasite membranes to such highly charged compounds. However, we found that simple chemical supplementation of the media with D-mannose was sufficient to rescue growth when HAD5 expression is reduced. This indicates that the primary mechanism of death in these parasites is due to defects in mannose metabolism (Fig. 2D, S2B). Notably, while artificially elevated concentrations of D-mannose completely rescued parasite growth, a physiologically relevant concentration of 50 µM [equivalent to that of human serum(47)] did not provide a statistically significant rescue of parasite growth.

### HAD5 is required for parasite egress and invasion

GPI anchors, which contain mannose(16, 18, 31), are required for egress of the malaria parasite from the infected host cell, as well as reinvasion of merozoites(46, 48–55). Because of its essential role in mannose metabolism, we hypothesized that HAD5 may be required for efficient egress and invasion. To evaluate this possibility, we washed out aTc at the beginning of the life cycle in synchronized parasites. Parasites grown in -aTc conditions over the course of one life cycle developed morphologically normally through the majority of life cycle stages, including the development of multi-nucleated schizonts. Whereas +aTc conditions allowed parasites to continue into the next life cycle and form newly reinvaded “ring”-stage parasites, -aTc parasites were arrested in late schizogony (Fig. 2E), which suggests a defect in parasite egress when HAD5 expression is knocked down. Notably, when HAD5 expression is reduced, parasites retain normal mature schizont architecture by transmission electron microscopy, indicating that neither gross developmental defects nor structural aberrations prevent parasites from egressing (Fig. S3).

To examine whether HAD5 is required for invasion, as well as egress, we examined the reinvasion capacity of HAD5^KD^ parasites. Segmented schizont-enriched cultures of HAD5^KD^ ±aTc were mechanically lysed and the freed merozoites were allowed to reinvade fresh RBCs. Under these conditions, knockdown parasites were unable to reinvade new host cells (Fig. S4). Thus, the knockdown of HAD5 confers defects to parasite biology that prevent both egress and reinvasion of the parasites.

### HAD5 knockdown disrupts GPI anchor synthesis

Mannose metabolism is linked to parasite egress through biosynthesis of GPI anchors(56). In *P. falciparum,* GPI anchors are synthesized through addition of one glucosamine (GlcN) and three to four mannose residues to a phosphatidylinositol backbone. These mannose residues are derived from the product of phosphomannomutase, M1P, which is converted to GDP-mannose and subsequently to dolichol-phosphate mannose, the direct mannose donor to GPI anchors (Fig. 3A)(57). Several GPI-APs contribute to egress and invasion of parasites(16, 19, 21). We therefore hypothesized that reduced HAD5 expression leads to loss of phosphomannomutase activity and causes parasite death by disruption of GPI anchor biosynthesis. To directly evaluate the effect of HAD5 knockdown on GPI anchor biosynthesis, we labeled mid- to late-trophozoite parasites with [^3^H]-GlcN and extracted GPI precursors as previously described(18, 56, 58, 59). HAD5^KD^ parasites grown in +aTc conditions had the expected repertoire of GPI anchor precursors (Fig. 3B, 3C). A variety of precursors are observed, with earlier, less polar species (with fewer mannose groups) migrating further than more polar, highly-mannosylated species. When HAD5 expression is reduced, there is a relative accumulation of the earlier precursors, as well as a reduced production of fully mature, highly mannosylated precursors (Fig. 3B), indicating a defect in GPI anchor biosynthesis. In particular, there was a significant reduction of the highly polar band 9 and significant buildup of less polar band 4 when HAD5 expression was reduced (Fig. 3C). Intriguingly, despite the substantial knockdown of HAD5 and the complete loss of growth in these parasites, the abundance of many mannosylated precursors is unchanged and highly mannosylated GPI precursors are still observed, suggesting that this biosynthetic pathway is not completely ablated.

**Figure 3.**
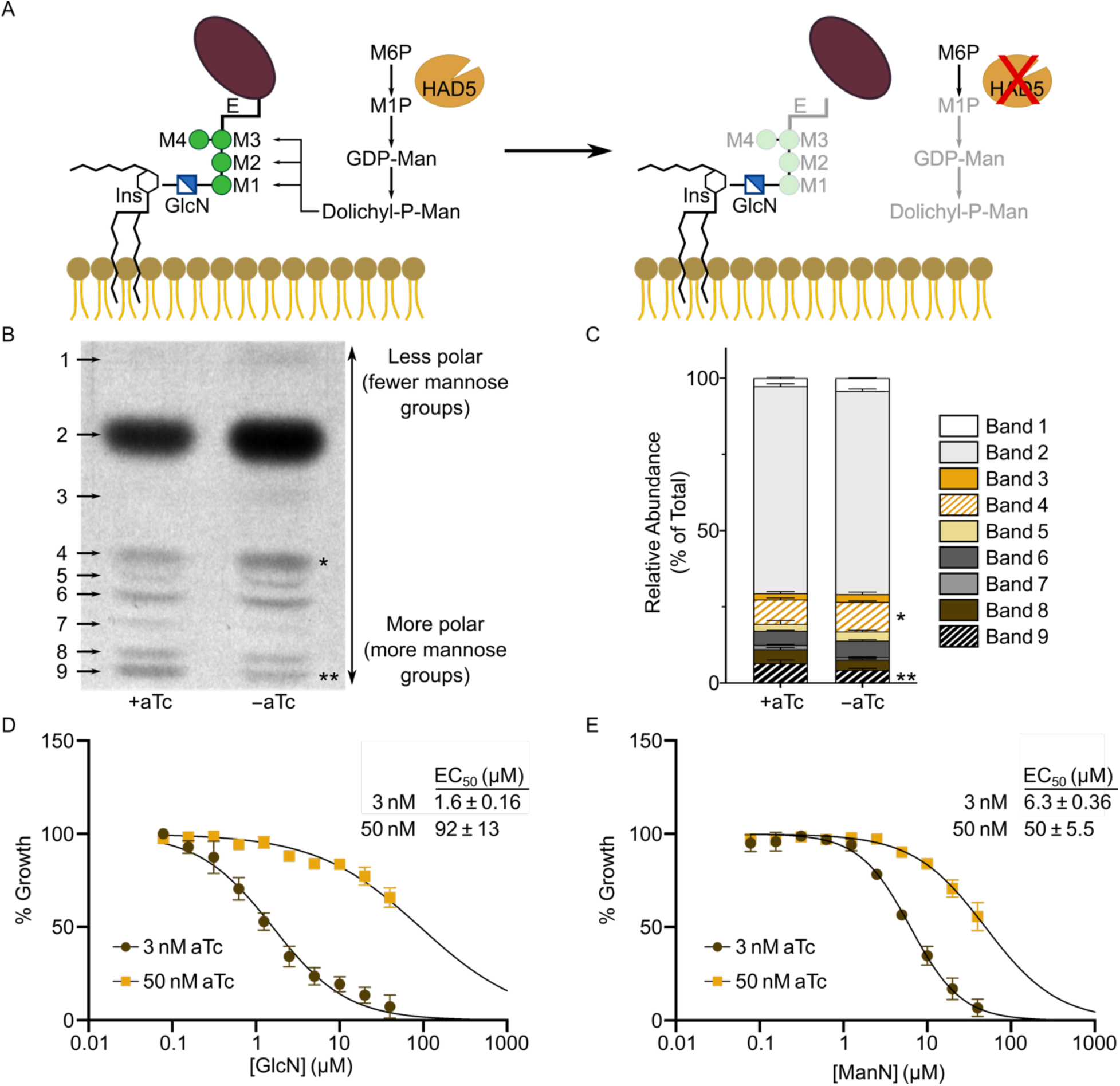
Knockdown of HAD5 disrupts GPI anchor biosynthesis. **(A)** Model of the predicted effect that knocking down HAD5 will have on GPI anchor precursor synthesis and subsequent anchoring of GPI-APs. Abbreviations: Ins, inositol; GlcN, glucosamine; M, mannose; E, ethanolamine; M6P, mannose 6-phosphate; M1P, mannose 1-phosphate. **(B)** Representative autoradiography film of GPI-anchor-precursors from [^3^H]GlcN-labeled parasites. Bands toward the top migrated farthest on a silica TLC plate, indicating they are less polar and less mannosylated. **(C)** The radiographic signal was quantified and is represented as proportions of total signal. Band numbers indicate the corresponding band from **A**. Shown are the mean and SEM of three independent experiments, analyzed by ordinary two-way ANOVA with Fisher’s LSD test. \**p=*0.038, \*\**p*=0.008. **(D,E)** Dose response curve of parasite growth in the presence of glucosamine (GlcN; **D**) or mannosamine (ManN; **E**). Data represent the means and SEM of three independent experiments with technical replicates.

To confirm the role of HAD5 in GPI biosynthesis, we deployed two established chemical tools, mannosamine (ManN) and GlcN, which impair growth of *P. falciparum* through inhibition of GPI anchor biosynthesis(60, 61). We expected that if HAD5 knockdown disrupts GPI anchor biosynthesis, parasites should be hypersensitized to ManN and GlcN treatment. Indeed, when expression of HAD5 is reduced with an intermediate concentration of aTc (3 nM) that still permits modest asexual growth, knockdown parasites yielded a marked shift in half-maximal effective concentration (EC_50_) (Fig. 3D, 3E). These results further implicate HAD5 in the production of GPI anchors in *P. falciparum*.

### HAD5-dependent GPI anchor synthesis enables proper anchoring of MSP1

Reduced GPI anchor biosynthesis in malaria parasites is expected to impact the localization and function of a number of essential GPI-anchored parasite proteins. While several GPI-anchored proteins have been characterized in *P. falciparum* intraerythrocytic stages, the most abundant is MSP1(16, 17). MSP1 must be targeted and anchored through GPIs and proteolytically processed in order for schizont-stage parasites to egress from the erythrocyte, and the MSP1 complex is also critical for binding and reinvading new red blood cells(19, 62). For this reason, we investigated whether HAD5-dependent GPI anchor synthesis is required for localization and anchoring of MSP1. We expected that, when the pathway is intact, MSP1 is successfully anchored to the parasite plasma membrane. In contrast, when HAD5 expression is knocked down, GPI anchors will fail to fully incorporate mannose and GPI-anchored proteins, including MSP1, will remain untethered to the membrane (Fig. 3A). To evaluate this effect, we used immunofluorescence to detect the localization of MSP1. When schizonts grown in ±aTc conditions were mechanically lysed and the resultant merozoites were imaged, there was a modest but significant decrease in MSP1 signal surrounding the daughter merozoites when HAD5 expression is reduced (Fig. 4A, 4B), indicating that MSP1 membrane attachment is diminished, causing it to diffuse away from the cell.

**Figure 4.**
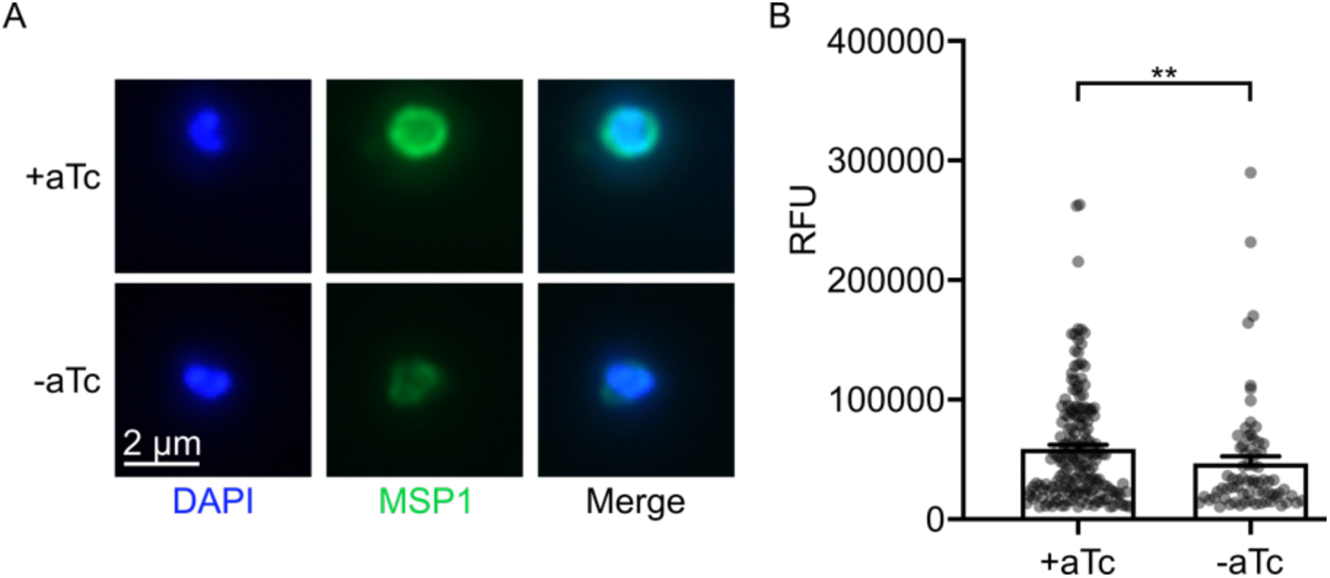
Knockdown of HAD5 diminishes membrane anchoring of the egress and invasion protein MSP1. **(A)** Representative immunofluorescent images of mechanically freed merozoites that were grown in ±aTc conditions and schizont enriched by E64 treatment. **(B)** Quantification of MSP1 signal from **A**. Data points represent three independent experiments, each with >25 observed merozoites, removing those under a threshold of 10,000 RFU. Bar graphs represent the mean + SEM of all data points. Statistics were performed by Mann-Whitney test. \*\**p*=0.008.

To independently confirm this finding, we partitioned lysate from early schizont-stage parasites into membranous and soluble fractions(63). Whole lysate and fractions were assessed by immunoblotting for MSP1, and the relative proportion of MSP1 in the membrane fraction compared to lysate was calculated. In +aTc conditions, a large proportion of MSP1 separates with the detergent-enriched phase, with some full-length MSP1 in the soluble phase. Upon HAD5 knockdown, there was a subtle reduction in the proportion of cellular MSP1 bound to the membrane (Fig. S5). This difference did not reach significance; however, substantial biological variability in MSP1 expression is present within merozoite populations, as indicated in Figure 4B, and post-hoc power analysis of this data indicates that we were underpowered to observe a significant difference (Power = 0.290)(64). Together with our analysis of GPI anchor biosynthesis in these parasites (Fig. 3), these data confirm some functional preservation of GPI biosynthesis when HAD5 expression is reduced.

### Disruption of mannose metabolism by HAD5 knockdown does not affect fosmidomycin sensitivity in parasites

HAD5 and other phosphomannomutases are part of the haloacid dehalogenase (HAD) superfamily of proteins (Interpro: IPR023214), a ubiquitous family of enzymes that primarily conduct phosphatase and phosphotransferase reactions(65). HAD5 is a member of subfamily IIB (IPR006379) of the HAD superfamily and has substantial sequence homology to other *P. falciparum* HAD proteins in this subfamily (HAD1 and HAD2) and the related subfamily IIA (IPR006357) protein, phosphoglycolate phosphatase (PGP)(66). One notable commonality between these subfamily II *P. falciparum* HAD proteins is their effect on the parasite’s sensitivity to the antimalarial fosmidomycin (FSM), which is a well-validated inhibitor of the apicoplast methylerythritol phosphate pathway of isoprenoid biosynthesis(10). Mutations in either HAD1(67) or HAD2(68) render parasites resistant to FSM, and PGP-knockout parasites are hypersensitive to FSM(69). To examine whether HAD5 also plays a role in FSM susceptibility, we assessed the FSM dose-response of parasites grown in saturating or intermediate aTc conditions. Unlike its close homologs, HAD5 knockdown had no effect on the FSM EC_50_ of parasites (Fig. S6).

### HAD5 is distinct from human PMMs and can be specifically inhibited

Previous studies have investigated egress and invasion as a promising target for antimalarial drug discovery, suggesting that HAD5 may likewise be of interest for antimalarial drug discovery(46, 48–51); however, phosphomannomutases are found widely throughout nature, including two genes in the human genome, *Hs*PMM1 and *Hs*PMM2(70, 71). We therefore evaluated the potential for selective inhibition of *P. falciparum* HAD5 over human PMM1 and PMM2 (Fig. S7A). Previous work has successfully demonstrated the use of substituted ketoheptoses and other phosphosugar analogues as inhibitors of microbial phosphomannomutases(72, 73), which we sought to replicate with HAD5. Using a panel of 11 phosphosugar analogues (Fig. S8A), we screened each compound for its ability to inhibit recombinant HAD5, PMM1, and PMM2. The majority of the compounds had negligible inhibition against all three enzymes (Fig. S8). Compound D9, however, inhibits purified recombinant HAD5 with a half-maximal inhibitory concentration of 79 ± 2.6 µM, several-fold more potently than the inhibition of either PMM1 or PMM2 (Fig. 5A). Moreover, we find dramatic time-dependent effects on the ability of D9 to inhibit HAD5, as pre-incubating HAD5 with D9 prior to assaying activity substantially increased D9 potency, such that a 60-minute pre-incubation yielded HAD5 activity of only 4.5% of a vehicle-treated control (Fig. 5B). This effect was not seen for *Hs*PMM1, demonstrating that the potential to specifically inhibit HAD5 may be even greater under ideal binding conditions. As expected given its poor drug-like characteristics (and likely inadequate cellular permeability), compound D9 did not impair the growth of asexual *P.* falciparum at concentrations up to 100 µM (Fig. S9A). The selective inhibition of HAD5 by D9 is an important proof-of-concept that distinct structural features of HAD5 may be harnessed for parasite-specific inhibitor development.

**Figure 5.**
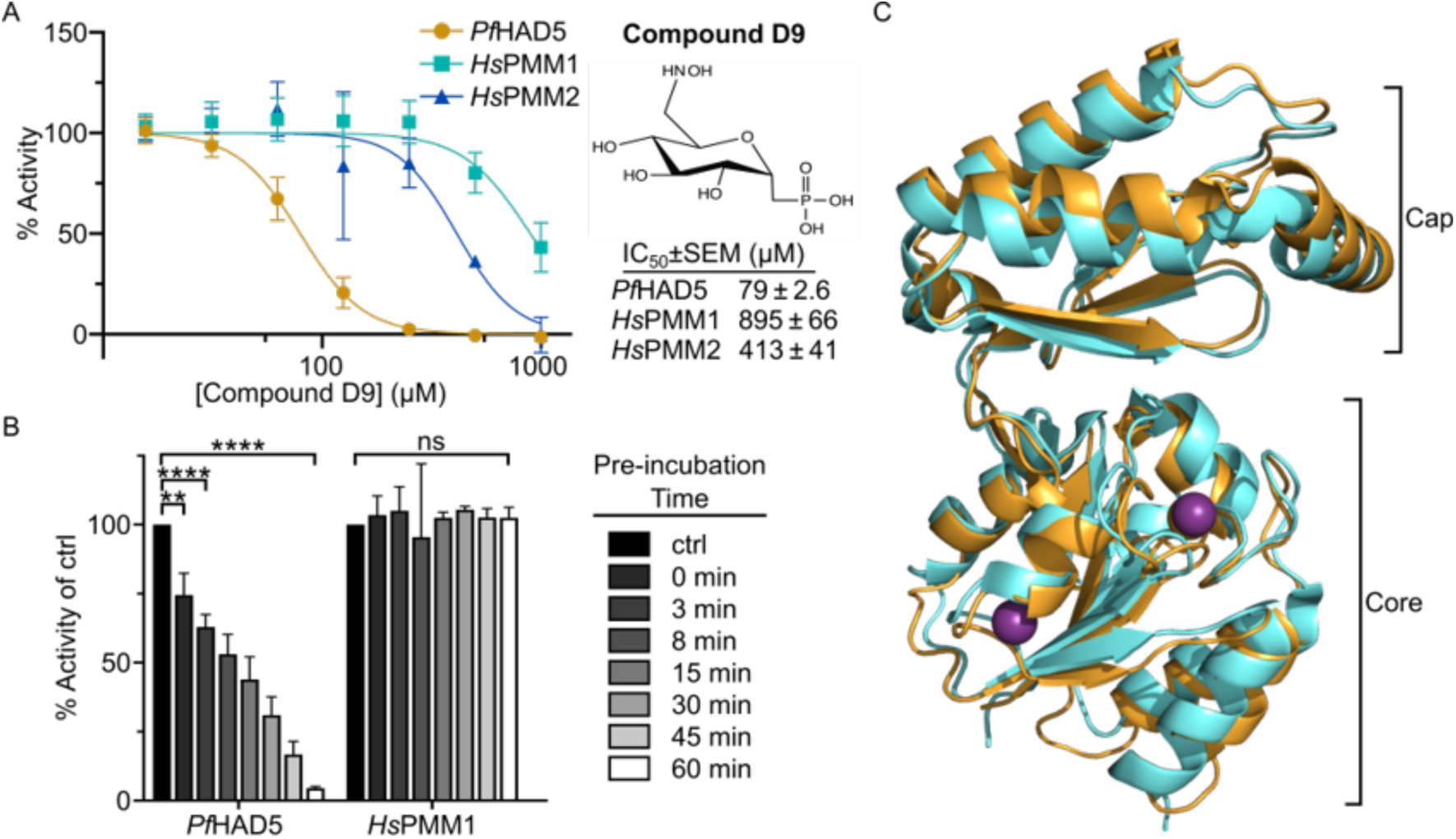
HAD5 is sufficiently distinct from human PMMs to be specifically inhibited. **(A)** Dose response curve of compound D9 against recombinant *Pf*HAD5, *Hs*PMM1, and *Hs*PMM2. Data represents the mean ± SEM of three independent experiments, each with technical replicates. **(B)** Activity of *Pf*HAD5 or *Hs*PMM1 was assayed after preincubating enzymes for the given time with 416 µM of D9 prior to adding the preincubation to the reaction mix (final [D9] = 50µM). As a control (ctrl), enzymes were incubated with an equal volume of water for 60 minutes. Statistics were performed with an ordinary Two-way ANOVA, using Dunnett’s test for multiple comparisons. \*\**p=*0.0046 \*\*\*\**p* < 0.0001, ns = not significant. **(C)** 3.5 Å resolution crystal structure of PfHAD5 (orange; PDB 7MYV) aligned to HsPMM1 (cyan; PDB 2FUC) with Mg^2+^ ions (purple). Indicated are the Cap and Core domains typical of HAD enzymes.

To uncover the structural basis for HAD5-specific inhibition, we solved the crystal structure of HAD5 to 3.5 Å resolution. Like other known phosphomannomutases and members of the HAD superfamily(65, 74), HAD5 comprises two domains with an overall fold similar to that of *Hs*PMM1 (Fig. 5C, Table S1). There were no large structural differences between the human and parasite enzymes, so we next evaluated the substrate-binding pocket by computationally docking D9 onto the HAD5 crystal structure (Fig. S9B). Although the overall binding pockets were similar and the amino acid sequence within the pocket are highly conserved, subtle differences in side-chain positioning may account for the parasite specificity of D9. Further analysis of the binding pocket is restricted by the current resolution of 3.5Å. Furthermore, comparing D9 to the inactive compounds D7 and D11, the singular difference is the presence of a hydroxyaminomethyl group on carbon 6 of compound D9, whereas an aminomethyl group or hydroxymethyl group are present on D7 and D11, respectively (Figure S8A), a difference that may explain the disparity in potency and specificity.

## DISCUSSION

We report here the lethal knockdown and biochemical characterization of the phosphomannomutase in *P. falciparum*, HAD5. Loss of HAD5 leads to growth arrest in asexually replicating parasites, marked by defects in egress and reinvasion. This growth defect can be rescued by media supplementation with D-mannose, indicating that disruption of mannose metabolism is the primary mechanism of death in HAD5^KD^ parasites; however, a physiologically relevant concentration of 50 µM D-mannose is unable to significantly rescue growth, bolstering the case for this pathway as a therapeutic target. We further report the specific inhibition of HAD5 enzymatic activity compared to orthologous human phosphomannomutases by the hexose-phosphate analogue, compound D9, highlighting the potential for specific therapeutics to be developed to HAD5.

Unexpectedly, despite the dramatic decrease in HAD5 protein levels and the compelling loss of growth upon HAD5 knockdown, GPI anchor synthesis and MSP1 anchoring to merozoite surfaces are only modestly impacted. Residual HAD5 activity may be present after knockdown and sufficient for generating detectable GPI precursors and subsequent tethering of some GPI-APs, including MSP1. Alternatively, *P. falciparum* expresses two annotated phosphoglucomutases (PGMs) (PF3D7_1012500 and PF3D7_0413500)(75) that could potentially catalyze phosphomannomutase activity as well, indicating some functional redundancy to HAD5. Residual HAD5 or functional redundancy by PGMs could also explain our observation that D-mannose rescues HAD5 knockdown. Intracellular mannose is likely converted to M6P by hexokinase, and thus some phosphomannomutase activity would still be required for successful rescue of the GPI-anchor pathway. In either case, while residual phosphomannomutase activity present in HAD5^KD^ may not eliminate GPI synthesis entirely, it is insufficient to sustain parasite growth. This speaks to an exquisite sensitivity of malaria parasite cells to disruption in this pathway, whereby minor perturbations in GPI anchor synthesis nonetheless completely interrupt parasite growth, highlighting the promise of this pathway as a therapeutic target.

In addition, we find that HAD5^KD^ parasites have a complete cell cycle arrest, although others have found that untethering MSP1 from the membrane by removing its GPI-anchoring C-terminus still allows for minimal parasite growth(19). We expect that this discrepancy is due to the role of HAD5 in function of all GPI-anchored parasite proteins, not solely MSP1. These other GPI-APs include related MSPs, RAMA, and 6-cysteine proteins, many of which are refractory to deletion and likely essential(16, 21, 35, 76). With the incomplete loss of GPI synthesis, it may be that each one of these GPI-APs, including MSP1, are only relatively de-anchored, but the modest reduction in this post-translational modification across multiple cellular proteins works in concert to cause parasite growth arrest.

We therefore propose that HAD5, as an upstream member of the GPI biosynthesis, has great potential as an antimalarial target. We expect that HAD5 inhibition will have broad downstream effects on parasite biology across several life cycle stages. That compound D9 has markedly improved potency against malaria HAD5 compared to orthologous human enzymes provides key proof-of-concept for ongoing development of specific HAD5-directed antimalarial therapeutics. While compound D9 has limited antimalarial efficacy, this is likely due to its charged phosphonate, expected to have poor cellular penetration. This liability may be improved through a variety of medicinal chemistry strategies, including the addition of pro-drug moieties to mask this charge, a strategy that has been highly effective for other phosphonate antimalarials in development (77–79). Finally, the crystal structure of HAD5 is likely to be valuable to ongoing efforts to develop more potent and specific HAD5 inhibitors as antimalarials.

This study also adds to the growing literature on HAD-like proteins in *P. falciparum*. Three related HAD proteins each independently modulate parasite sensitivity to the isoprenoid biosynthesis inhibitor FSM, which prompts the question of whether this effect would be similarly seen with other HAD proteins(67–69). Alternatively, the non-mevalonate isoprenoid biosynthesis pathway may be particularly sensitive to cellular metabolic perturbations. We find that HAD5 serves as an interesting counterexample. HAD5 knockdown yields no changes to FSM sensitivity, providing evidence of a HAD protein and a metabolic perturbation that does not impact the sensitivity of parasites to inhibition of isoprenoid metabolism.

Finally, we note that HAD5 and the GPI anchor biosynthesis pathway are expressed throughout the parasite’s life cycle. Several GPI-anchored proteins are expressed in gamete- and oocyst-stage parasites(80, 81), and CSP, the target of the RTS,S malaria vaccine and the more recent R21 vaccine candidate(26–29), is a GPI-anchored protein expressed on the surface of sporozoites that facilitates sporozoite development(30) and targets sporozoites to the liver(82). Furthermore, sexual-stage parasites harbor additional fates of mannose 1-phosphate, including *C*-mannosylation(83, 84) and *O*-fucosylation(85, 86), suggesting that HAD5 will play a critical role in parasite biology across several life stages. Hence, HAD5 serves an essential role in parasite metabolism and shows promise as a specific therapeutic target. The inhibition of HAD5 not only has potential to treat intraerythrocytic *P. falciparum* infection, but may also serve to block transmission.

## EXPERIMENTAL PROCEDURES

### Parasite Strains and Culturing

Unless otherwise indicated, parasites were maintained at 37°C in 5% O_2_, 5% CO_2_, 90% N_2_ in a 2% suspension of human red blood cells in RPMI medium (Gibco) modified with the addition of 30 mM NaHCO_3_, 11 mM glucose, 5 mM HEPES, 1 mM sodium pyruvate, 110 µM hypoxanthine, 10 µg/mL gentamicin, (Sigma) and 2.5 g/L AlbuMAX I (Gibco). Deidentified RBCs were obtained from the Barnes-Jewish Hospital blood bank (St. Louis, MO), St. Louis Children’s Hospital blood bank (St. Louis, MO), and American Red Cross Blood Services (St. Louis, MO). The HAD5 conditional knockdown strain, “HAD5^KD^”, was generated by transfecting NF54^attB^ parasites(87) using previously described methods(45, 46). Resultant parasites were maintained in the presence of anhydrotetracycline (aTc; Cayman Chemicals) in DMSO at 500 nM unless otherwise specified. Parasites were synchronized with a combination of 5% sorbitol (Fisher Bioreagents) and 1.5 µM Compound 1 (MedChemExpress)(88, 89)

### Plasmodium growth measurement

Parasitemia in daily growth assays was measured via flow cytometry by incubating 10 µL of parasite culture with 190 µL of 0.4 µg/mL acridine orange (Invitrogen) in PBS for 1 minute. Stained parasites were analyzed on a BD FACS Canto flow cytometer gating on DNA and RNA-bound dye signal using FITC and PerCP-Cy5.5 filters, respectively. 50,000 events were recorded for each sample.

### Merozoite reinvasion

Merozoite reinvasion was assessed as previously described(90). Parasites were synchronized, grown in ±aTc, and treated with 10 µM epoxysuccinyl-L-leucylamido(4-guanidino)butane (E64) (Sigma) for 8 hours to stall them in late schizont stages. E64 was washed out, and cultures were passed through a 1.2 µm filter to lyse the schizonts. Lysates were incubated with fresh red blood cells for 1 hour to allow reinvasion before cultures were washed again to remove debris. Parasitemia was assessed 24 hours post-reinvasion by acridine orange staining and flow cytometry. Reinvasion efficiency was assessed by normalizing resultant parasitemia to the measured parasitemia prior to lysis.

### Light and Fluorescent Microscopy

Parasite development was monitored by thin smear of synchronized parasites that were dyed with modified Giemsa Stain (Sigma). For fluorescent microscopy, E64-treated schizont stage parasites were mechanically lysed by passing them through a 1.2 µm filter(90). Lysates were added to poly-lysine-coated glass slides, fixed in 4% paraformaldehyde, 0.0075%glutaraldehyde in PBS, and permeabilized in 0.1% TritonX-100. Immunofluorescence was performed using mouse anti-MSP1 monoclonal antibody (Novus Biologicals) and goat-anti-mouse Alexafluor 488 secondary antibody (Life Technologies). Samples were then preserved with ProLong Gold antifade reagent with DAPI (Life Technologies).

All immunofluorescence and bright field images were taken using a Zeiss AxioObserver D1 inverted microscope (Carl Zeiss Inc.), equipped with a Axiocam 503 color camera, at the Washington University Molecular Microbiology Imaging facility. Images were acquired with a Plan-Apochromat 100x (NA 1.4) objective using the ZEN 2.3 pro (blue edition) software. Fluorescent signal was quantified using the ImageStudioLite software from LI-COR.

### Transmission Electron Microscopy

For ultrastructural analyses, highly synchronized infected RBCs were fixed in 2% paraformaldehyde/2.5% glutaraldehyde (Polysciences Inc., Warrington, PA) in 100 mM sodium cacodylate buffer, pH 7.2 for 1 hr at room temperature. Samples were washed in sodium cacodylate buffer and postfixed in 1% osmium tetroxide (Polysciences Inc.) for 1 hr at room temperature. Samples were then rinsed in dH_2_0, dehydrated in a graded series of ethanol, and embedded in Eponate 12 resin (Ted Pella Inc., Redding, CA). Sections of 95 nm were cut with a Leica Ultracut UCT ultramicrotome (Leica Microsystems Inc., Bannockburn, IL), stained with uranyl acetate and lead citrate, and viewed on a JEOL 1200 EX transmission electron microscope (JEOL USA Inc., Peabody, MA) equipped with an AMT 8 megapixel digital camera and AMT Image Capture Engine V602 software (Advanced Microscopy Techniques, Woburn, MA).

### Cloning

The coding sequence of HAD5 was cloned from cDNA of 3D7 parasites using primers P1 and P2 (Table S2) and cloned into the BG1861 vector(91), which introduces an N-terminal 6xHis-tag, by ligation independent cloning (LIC). This coding sequence was subsequently cut-and-pasted into a pET28a vector with NdeI and BamHI-HF (NEB), followed by ligation with NEB Quick Ligase using manufacturer’s protocols. The coding sequence did not match published reference sequence of PF3D7_1017400, as an adenosine-to-guanosine mutation yielded an Asn-to-Ser substitution at residue 100. To revert this sequence to the reference sequence, primer P3 (Table S2) was used in the QuikChange Multisite-directed mutagenesis kit (Agilent), and the resulting plasmid was transformed into XL10 Gold ultracompetent *Escherichia coli* cells. The HAD5^D11A^ allele was generated from the WT plasmid, again using the QuikChange multisite-directed mutagenesis kit and primer P4 (Table S2). *Homo sapiens* PMM1 and PMM2 coding sequences were identified from UniProt, codon-optimized for *E. coli*, and synthesized by Integrated DNA Technologies (Table S3). These gene blocks were amplified and extended using primers P5+P6 (PMM1) and P7+P6 (PMM2) and PrimeSTAR GXL DNA polymerase (Takara)(Table S2). These sequences were then cloned into the pET28a plasmid by Gibson assembly at NdeI and BamHI-HF cut sites, and plasmids were transformed into XL10 Gold ultracompetent *E. coli* cells.

*E. coli* mannose-1-phosphate guanylyl transferase (ManC) was cloned for use in phosphomannomutase assays. The coding sequence of *Ec*ManC was identified from UniProt and a gene block of the sequence was ordered from Integrated DNA Technologies (Table S3). This sequence was cloned by LIC into a BG1861 vector that had been modified with a starting KFS sequence downstream of the 6xHis-tag to enhance protein expression(92), and the resulting plasmid was transformed into Stellar competent cells (Takara).

### Recombinant Protein Expression & Purification

To generate recombinant protein for enzyme assays and crystallography, the pET28a plasmids were transformed into BL21 Gold (DE3) competent *E. coli* cells (Agilent). Cells were grown to optical density at 600nm (OD_600_) of 0.6-0.8 in Terrific Broth medium (24 g/L yeast extract, 20 g/L tryptone, 4 mL/L glycerol, 17 mM KH_2_PO_4_, 72 mM K_2_HPO_4_) shaking at 37°C, and then induced for approximately 18 hours with 1mM isopropyl-beta-D-thiogalactoside (IPTG)at 16°C. Cells were collected by centrifugation and resuspended in lysis buffer containing 25 mM Tris HCl (pH 7.5), 250 mM NaCl, 1 mM MgCl_2_, 10% glycerol, 20 mM imidazole, and 100 µM phenylmethylsulfonyl fluoride (PMSF) and lysed by sonication. Lysate was centrifuged at 18,000xg for 1 hr and the supernatant containing soluble protein was bound to nickel agarose beads (Gold Biotechnology), washed with buffer containing 25mM Tris HCl (pH 7.5), 250 mM NaCl, 1 mM MgCl_2_, 10% glycerol, 20 mM imidazole, and eluted with 10 mL of buffer containing 25mM Tris HCl (pH 7.5), 250mM NaCl, 1mM MgCl_2_, 10% glycerol, and 320mM imidazole. Eluate was dialyzed overnight at 4°C in the presence of 30 units of thrombin protease (Sigma) into buffer containing 25 mM Tris HCl (pH 7.5), 250mM NaCl, 1 mM MgCl_2_, 10% glycerol. The dialyzed, thrombin-cleaved mixture was then run over nickel agarose beads and benzamidine-sepharose beads (GE Healthsciences) to remove the cleaved His-tag and thrombin. The flow through protein was collected and further purified by size-exclusion chromatography using a HiLoad 16/60 Superdex-200 column (GE Healthsciences), equilibrated in dialysis buffer. Elution fractions containing the protein of interest were identified by UV absorbance and SDS-PAGE, pooled, and concentrated to 6 mg/mL. Protein solutions were flash frozen in liquid nitrogen and stored at -80°C (Figure 1B, S7A).

*Ec*ManC for enzyme activity assays was expressed by cloning the BG1861:ManC vector into BL21(DE3) pLysS *E. coli* cells (Life Technologies). Cells were grown in LB broth to an OD_600_ of 0.7-0.8 and induced with 1mM IPTG for 2hr at 37°C. Cells were harvested and protein was purified as described for other proteins above. However, as this construct lacked the thrombin-cleavable site from the pET28a vector, the elution from nickel beads was directly run over the size exclusion column, then pooled and concentrated (Figure S7B).

### HAD5 activity assays

Phosphomannomutase activity of HAD5 was measured using a linked enzyme scheme (Fig. S1A), modified from the EnzChek phosphate release kit (ThermoFisher). 200 µM of 2-amino-6-mercapto-7-methyl-purine riboside (MESG) and 1x EnzChek reaction buffer (50 mM Tris HCl, 1 mM MgCl_2_, pH 7.5, containing 100 µM sodium azide) were incubated with 1 U/mL Purine Nucleoside Phosphorylase (PNP), 1 U/mL pyrophosphatase, 125 µM GTP, 52 µg/mL recombinant purified *E. coli* ManC, and 1mM mannose-6-phosphate. 12.5 µM glucose-1,6-bisphosphate was also added to demonstrate its activating properties on HAD5. All reagents were obtained from Sigma-Aldrich. The reaction was started by adding 50 ng recombinant HAD5 (WT or D11A), which is within the linear range of the assay (Figure S1B). Reactions took place in 40 µL, and the production of nucleotide by PNP was measured at 360nM.

Phosphoglucomutase assays (Fig. S1C) were developed by incubating 1mM glucose-1-phosphate, 0.75 mM NADP^+^, 2.5 U/mL glucose-6-phosphate dehydrogenase, and 10 µM glucose-1,6-bisphosphate in 50 µL reactions containing 25 mM NaCl, 25 mM Tris HCl pH 7.5, and 1 mM MgCl_2_. All reagents were obtained from Sigma-Aldrich. Again, 50 ng of HAD5 was added to start the reaction (Fig. S1D), and the production of reduced NADPH was measured by absorbance at 340 nm.

All reactions took place at room temperature in clear CoStar 96-well half-area plates and absorbances were measured by a Perkin Elmer multimode Envision plate reader. For Michaelis-Menten kinetics of each assay, two-fold serial dilutions of substrate concentrations (either M6P or G1P, respectively) were made and all other components of the assays were kept constant.

### Inhibition assays of recombinant HAD5

Synthesis of compounds D1-D11 (Fig. S8A) have been described previously(72, 73), with the exception of compound D9, whose synthesis is described, and characterization data included, in the attached supporting information. To assess inhibition of HAD5, compounds D1-D11 were suspended in water and added to the phosphoglucomutase assay of HAD5 activity (the phosphomannomutase assay was not used, as cross inhibition was seen with downstream components of that assay, but not with the phosphoglucomutase assay; Fig S8D). 5 µL of water volume in the assay was replaced with 5 µL serial dilutions of analogs, with final concentrations ranging from 0 µM – 1 mM. The rate of product formation in each condition was used to determine the half maximal inhibitory concentration (IC_50_) for each inhibitor. In addition to HAD5, these assays were performed with recombinantly purified human PMM1 and PMM2. For these inhibition assays, 200 ng of HAD5 (139 nM), PMM1 (133 nM), or PMM2 (65 nM) was used to start the reaction.

Time-dependent inhibition was assessed by pre-incubating HAD5 or PMM1 in 416 µM D9 for time points up to 60 minutes, before adding the enzyme+inhibitor mixture to the full reaction mixture, achieving a final [D9] of 50 µM.

### Parasite growth inhibition assays

Dose-response inhibition experiments were performed on asynchronous parasites. Three experimental replicates were performed, with technical duplicates, for each experiment. GlcN and ManN inhibition experiments were performed by adding 2-fold dilutions of each compound in water, from 0 mM to 2 mM, to a 100 µL culture of parasites. 1 µM of chloroquine was used as a positive control. Inhibition of cultured parasites by compound D9 was assessed using 100 µM D9 or equivalent volume of water. Parasite growth in these experiments was assessed by acridine-orange staining and flow cytometry measurement of parasitemia 72 hours after treatment.

Dose-response of fosmidomycin (FSM) was assessed using concentrations up to 50 µM FSM in water, and growth was measured after 72 hour treatment on a PerkinElmer multimode Envision plate reader by Quant-iT PicoGreen dsDNA reagent (Thermofisher) staining.

### Western blotting

To verify tagging and knockdown of protein, cultures of HAD5^KD^ were grown for 24 hours in ±aTc conditions, then RBCs were lysed with cold PBS + 0.1% saponin. Samples were centrifuged to pellet parasites and remove excess hemoglobin, then parasites were lysed in RIPA (50 mM Tris, pH 7.4; 150 mM NaCl; 0.1% SDS; 1% Triton X-100; 0.5% DOC) plus HALT-Protease Inhibitor Cocktail, EDTA-free (Thermo Fisher). Lysates were centrifuged at high speed to pellet and remove hemozoin. Cleared lysates were then diluted in SDS sample buffer (10% SDS, 0.5 M DTT, 2.5 mg/mL bromophenol blue, 30% 1 M Tris pH 6.8, 50% glycerol) and boiled for 5 min. Lysates were separated by SDS-PAGE, then transferred to 0.45 μm nitrocellulose membrane (Thermo Scientific). Membranes were blocked in PBS + 3% bovine serum albumin, then probed with primary antibodies: mouse anti-FLAG (M2, Sigma), rabbit anti-HSP70 (Agrisera). Membranes were washed in PBS + 0.5% Tween 20, then probed with secondary antibodies goat anti rabbit IRDye 800CW 1:20,000 (LI-COR) and donkey anti-mouse IRDye 680LT 1:20,000 (LI-COR). Membranes were then washed in PBS + 0.1% Tween 20 and imaged on a LI-COR Odyssey platform.

### Autoradiography and GPI anchor quantification

GPI anchor autoradiography was performed as previously described(18, 56, 58). Briefly, 33-39 hr old parasites that had been grown in ±aTc conditions were washed and resuspended in glucose-free media (RPMI R1383 + 20 mM D-fructose, 25 mM HEPES, 21 mM NaHCO_3_, 0.37 mM hypoxanthine, 11 mM glutathione, 5 g/L Albumax I, and 10 µg/mL gentamicin). 150 µCi of 40 Ci/mmol [^3^H]GlcN (American Radiochemicals) was added to each 10 mL culture and incubated for 3 hours. Parasites were saponin lysed and glycolipids were extracted with chloroform:methanol:water (10:10:3), nitrogen evaporation, and n-butanol partitioning(59). Resultant glycolipids were run on TLC Silica gel 60 F_254_ plates (Millipore Sigma) using chloroform:methanol:water (10:10:3) as a solvent. TLC plates were exposed to autoradiography films (MidSci), which were developed after 1 week of exposure. Films were imaged on a BIO-RAD ChemiDoc MP imaging system, and signal was quantified by the ImageLab software from Bio-Rad.

### TritonX-114 Membrane Partitioning

Membrane partitioning was adapted from Doering et al.(63). Briefly, at approximately 40-44 hours post invasion, synchronized parasite cultures were magnetically purified (Miltenyi Biotec), and elution was centrifuged and resuspended in 1 mL ice-cold Tris-buffered saline (TBS) + 1x HALT protease inhibitor (PI) cocktail, EDTA-free (Thermo Fisher). 200 µL precondensed Triton X-114 was added to lyse parasites, and lysates were incubated on ice for 15 min. Lysates were centrifuged to remove debris. Then followed a series of 5 extractions with cold TBS+PI and precondensed TritonX-114, followed by warming to 37°C and centrifugation to separate phases. The lysate, detergent-enriched phase, and the soluble phase were analyzed by SDS-PAGE and Western blotted as described above using mouse anti-plasmepsin V 1:20(93) and rabbit anti-HAD1 1:10(67), or rabbit anti-MSP1 1:1000 MSP1(94). Detergent samples were diluted 5-fold in TBS prior to SDS-PAGE to avoid TritonX-114 interference with the gel.

### Protein Crystallography and Ligand Docking

Crystals of *P. falciparum* HAD5 were grown at 4°C using vapor diffusion in hanging drops comprised of 2 µL protein (6 mg/mL) and 2 µL crystallization buffer (0.1M bis -tris-propane pH 6.5, 0.1 M calcium acetate, 16% (w/v) PEG 8000, and 2% (v/v) benzamidine hydrochloride). Prior to data collection, crystals were flash frozen in mother liquor supplemented with 25% glycerol. All diffraction images were collected at 100K at beamline 19-ID of the Structural Biology Center at Argonne National Laboratory Advanced Photon Source. HKL3000 was used to index, integrate, and scale the data sets(95). Prior to phasing, a homology model of PfHAD5 was created using SWISS-MODEL(96) based on human alpha phosphomannomutase 1 (PDB: 6CFR(97), sequence identity 51%). Molecular replacement was performed using PHASER(98) and the homology model of PfHAD5 as a search model. COOT and PHENIX were used for iterative rounds of model building and refinement(99, 100). Data collection and refinement statistics are summarized in Table S1. Atomic coordinates and structure factors of PfHAD5 have been deposited in the RCSB Protein Data Bank (PDB: 7MYV).

The PfHAD5 structure was prepared for docking using AutoDock Tools 1.5.7(101). The three-dimensional structure of compound D9 was prepared using Avogadro(102), and was prepared for docking using AutoDock Tools 1.5.7. Docking was performed with AutoDock Vina using default search settings(103).

## ACKNOWLEDGMENTS

We thank Dr. Tamara Doering (Dept. of Microbiology, Washington University School of Medicine) for resources and technical assistance with GPI-anchor detection and membrane partitioning, Dr. Wandy Beatty (Molecular Microbiology Imaging Facility, Washington University School of Medicine) for electron microscopy and immunofluorescence assistance, and Dr. Akhil Vaidya (Dept. of Microbiology & Immunology, Drexel University College of Medicine) for *anti*-MSP1 antisera.

## SUPPORTING INFORMATION

This article contains supporting information.

## AUTHOR CONTRIBUTIONS

Provision and synthesis of compounds D1-D11 was by ES, JSZ, and DLJ. Recombinant protein synthesis and crystallography was performed by PMF and JJM. Crystal structure was solved and refined by PMF, JJM, and JMJ. All other experiments were performed by PMF. PMF, AROJ, and DEG were involved in experimental design, data interpretation, and the writing of this manuscript, with input from other authors.

## FUNDING AND ADDITIONAL INFORMATION

A.O.J. is supported by NIH/NIAID R01 AI103280, R21 AI123808, R21 AI130584, and R61 DH105594. AOJ is an Investigator in the Pathogenesis of Infectious Diseases (PATH) of the Burroughs Wellcome Fund. D.L.J is supported by NSERC (03893-2020) and CIHR (153332). The content is solely the responsibility of the authors and does not necessarily represent the official views of the National Institutes of Health.

## ABBREVIATIONS

aTc: anhydrotetracycline
CSP: circumsporozoite protein
E: ethanolamine
E64: epoxysuccinyl-L-leucylamido(4-guanidino)butane
FSM: fosmidomycin
G-1,6-P: glucose-1,6-bisphosphate
G1P: glucose 1-phosphate
G6P: glucose 6-phosphate
GlcN: glucosamine
GPI: glycosylphosphatidylinositol
GPI-AP: GPI-anchored protein
HAD: haloacid dehalogenase
Ins: inositol
LIC: ligation-independent cloning
M1P: mannose 1-phosphate
M6P: mannose 6-phosphate
Man: mannose
ManN: mannosamine
MSP: merozoite surface protein
PGM: phosphoglucomutase
PGP: phosphoglycolate phosphatase
PMM: phosphomannomutase
RAMA: rhoptry-associated membrane antigen

