## Supporting information for "Enzymatic and structural characterization of HAD5, an essential phosphomannomutase of malaria parasites"

**List of Materials**

Table S1: Summary of crystallographic statistics.
Table S2: Primers used for cloning.
Table S3: gBlock sequences used in cloning.
Figure S1: Enzymatic activity assays.
Figure S2: Chemical rescue of parasite growth.
Figure S3: Ultrastructures of HAD5$^{KD}$ schizonts by transmission electron micrograph.
Figure S4: HAD5 KD Parasites are deficient in reinvasion.
Figure S5: TritonX-114 partition of MSP1.
Figure S6: Knockdown of HAD5 has no effect on FSM sensitivity.
Figure S7: Purified recombinant proteins.
Figure S8: Structures and evaluation of compounds D1-D11.
Figure S9: Compound D9 does not inhibit parasites in culture.
Synthesis of phosphonate analogue of 6NHOH-G1CP (D9)

Scheme 1. Synthesis of phosphonate analogue of 6NHOH-G1CP (D9)
Figure S10: $^{1}$H NMR of compound **2**
Figure S11: $^{31}$P NMR of compound **2**
Figure S12: $^{13}$C NMR of compound **2**
Figure S13: $^{1}$H NMR of compound **3**
Figure S14: $^{31}$P NMR of compound **3**
Figure S15: $^{13}$C NMR of compound **3**
Figure S16: $^{1}$H NMR of compound **4**
Figure S17: $^{31}$P NMR of compound **4**
Figure S18: $^{1}$H NMR of compound **5**
Figure S19: $^{31}$P NMR of compound **5**
Figure S20: $^{13}$C NMR of compound **5**
Figure S21: $^{1}$H NMR of compound **6**
Figure S22: $^{31}$P NMR of compound **6**
Figure S23: $^{13}$C NMR of compound **6**
Figure S24: $^{1}$H NMR of compound **D9**
Figure S25: $^{31}$P NMR of compound **D9**
Figure S26: $^{13}$C NMR of compound **D9**

**Table S1. Summary of crystallographic statistics.**

| Crystal | PfHAD5 |
|---|---|
| Space group | $P6_522$ |
| Cell dimensions | *a*=*b*=161.3. Å, *c*=109.7 Å |
| Data Collection | |
| Wavelength | 0.979 Å |
| Resolution range (highest shell) | 40.3-3.50 Å (3.56-3.520 Å) |
| Reflections (total/unique) | 354,596 / 11,030 |
| Completeness (highest shell) | 99.9% (100%) |
| <I/□> (highest shell) | 18.5 (2.7) |
| $R_{sym}$ (highest shell) | 33.4% (143%) |
| Refinement | |
| $R_{crys}$ / $R_{free}$ | 25.4% / 31.0% |
| No. of protein atoms | 3,941 |
| No. of ligand atoms | 2 |
| R.m.s. deviation, bond lengths | 0.003 Å |
| R.m.s. deviation, bond angles | 0.74° |
| Avg. B-factor: protein, ligand | 92.7, 59.1 $Å^2$ |
| Stereochemistry: favored, allowed, outliers | 94.7, 4.5, 0.8 % |

**Table S2**: **Primers used for cloning**

| Primer Name | Sequence | Notes |
|---|---|---|
| P1 | CTCACCACCACCACCACCATATGAATAAGAAAAAAGGCAATATTTCTGT | For cloning HAD5 sequence from *P. falciparum* cDNA into BG1861 vector |
| P2 | ATCCTATCTTACTCACTTACAAGAAATTCTCTCTTAAAATTTTAAC | For cloning HAD5 sequence from *P. falciparum* cDNA into BG1861 vector |
| P3 | GAGAAGACCGATTAAAAAAATTGATAAATTATAGTTTAAAATATATTGCC | For reverting the HAD5 gene sequence to the reference sequence. |
| P4 | GGCAATATTTCTGTTTGCTGTAGATGGGACCC | For generating HAD5$^{D11A}$ mutant for recombinant protein. |
| P5 | GCCGCGCGGCAGCCATATGGCAGTTACA | Forward primer for cloning HsPMM1 gblock into pET28a vector. |
| P6 | CGGAGCTCGAATTCGGATCCTATCTTACTC | Reverse primer for cloning HsPMM1 and Hs PMM2 gblocks into pET28a vector. |
| P7 | CGGAGCTCGAATTCGGATCCTATCTTACTCACTTA | Forward primer for cloning HsPMM2 gblock into pET28a vector. |

**Table S3: gBlock sequences used in cloning**

| Gene Name | Sequence |
|---|---|
| *Hs*PMM1 | ATGGCAGTTACAGCCCAGGCAGCCCGTCGTAAGGAGCGTGTCTTATGTCTGT<br>TCGATGTAGACGGAACTCTGACCCCCGCACGTCAAAAAATCGACCCGGAAGT<br>TGCAGCTTTTTTGCAGAAGCTGCGTTCGCGCGTCCAGATCGGTGTAGTCGGC<br>GGATCAGATTACTGCAAAATCGCCGAGCAACTTGGAGATGGCGACGAAGTGA<br>TCGAGAAGTTTGACTACGTCTTCGCCGAGAATGGGACAGTTCAATACAAGCA<br>TGGGCGCTTATTGAGTAAGCAGACTATTCAGAACCATCTGGGGGAGGAGTTG<br>CTTCAAGATCTTATTAATTTTTGTTTATCCTATATGGCCTTACTTCGCCTGCCC<br>AAAAAGCGCGGTACTTTCATTGAGTTCCGTAACGGGATGCTGAACATCAGTC<br>CAATCGGTCGCTCATGCACTCTGGAGGAGCGTATCGAGTTTTCTGAACTTGA<br>CAAGAAAGAGAAAATTCGTGAGAAATTCGTCGAGGCGTTAAAAACGGAGTTT<br>GCAGGGAAGGGATTACGCTTTTCTCGCGGAGGCATGATTTCATTCGACGTGT<br>TTCCAGAAGGTTGGGACAAGCGCTACTGCTTGGACTCATTAGATCAAGATAG<br>CTTTGATACCATTCACTTTTTCGGGAACGAAACCTCGCCTGGGGGTAACGACT<br>TCGAGATCTTTGCGGACCCTCGTACGGTCGGGCACTCGGTAGTGAGCCCTC<br>AGGACACCGTGCAACGTTGTCGTGAGATTTTTTTCCCAGAGACGGCGCATGA<br>AGCGTAAGTGAGTAAGATAGGATCCGAATTCGAGCTCCG |
| *Hs*PMM2 | ATGGCGGCTCCGGGCCCAGCATTATGTTTATTTGACGTTGACGGAACCCTTA<br>CCGCACCGCGTCAAAAGATCACGAAGGAAATGGATGATTTTTTGCAGAAGTT<br>ACGTCAGAAGATCAAAATCGGGGTGGTCGGTGGTTCCGATTTTGAGAAAGTT<br>CAGGAGCAGCTTGGAAACGACGTGGTTGAGAAGTACGATTACGTCTTTCCGG<br>AAAATGGGTTGGTCGCGTATAAGGACGGTAAACTGCTTTGTCGTCAAAATATT<br>CAGTCCCATCTGGGCGAAGCCTTGATTCAAGATTTAATCAATTATTGCTTATC<br>CTATATCGCTAAGATCAAATTGCCCAAGAAACGCGGCACCTTTATTGAGTTTC<br>GTAATGGCATGTTGAACGTGTCCCCGATCGGACGTTCGTGTTCCCAGGAGGA<br>ACGCATTGAGTTCTATGAACTGGATAAAAAAGAAAATATCCGTCAAAAGTTCG<br>TTGCCGATCTTCGCAAGGAGTTCGCAGGCAAAGGTTTAACGTTCTCAATCGG<br>CGGTCAAATCTCTTTCGATGTGTTCCCAGACGGATGGGACAAACGTTACTGT<br>CTTCGCCATGTAGAAAATGATGGATATAAAACCATCTACTTTTTTGGGGACAA<br>AACAATGCCAGGAGGGAATGACCATGAAATTTTCACGGACCCCCGTACAATG<br>GGCTACTCAGTAACCGCACCGGAAGATACCCGTCGTATTTGCGAGCTGTTGT<br>TCTCTTAAGTGAGTAAG |

| | |
|---|---|
| *Ec*ManC | CTCACCACCACCACCACCATATGGCTCAAAGTAAACTTTACCCAGTGGTAATG<br>GCGGGGGGAAGTGGCTCTCGTCTTTGGCCTTTATCTCGCGTTCTTTATCCAA<br>AACAGTTCTTGTGCCTTAAGGGTGATTTGACAATGTTGCAAACAACGATCTGC<br>CGCCTGAATGGTGTCGAATGTGAGTCACCTGTGGTAATTTGCAATGAACAGC<br>ACCGTTTCATCGTAGCAGAACAACTGCGTCAGCTGAATAAATTAACGGAAAAT<br>ATTATCCTGGAGCCTGCTGGTCGCAACACGGCACCTGCAATCGCCTTGGCTG<br>CACTTGCGGCCAAGCGTCACTCACCAGAAAGTGACCCGCTTATGCTTGTCTT<br>GGCCGCCGATCATGTGATCGCAGATGAAGATGCTTTTCGTGCTGCTGTCCGT<br>AATGCGATGCCATATGCCGAAGCTGGAAAGTTAGTTACGTTTGGTATCGTGC<br>CGGATTTGCCGGAGACTGGATATGGTTATATCCGCCGTGGGGAGGTCAGCG<br>CTGGAGAACAAGACATGGTAGCCTTCGAGGTGGCACAATTTGTTGAAAAGCC<br>AAATCTTGAGACAGCACAAGCGTACGTGGCTAGTGGGGAGTATTACTGGAAC<br>TCTGGGATGTTTTTATTCCGTGCGGGGCGTTACCTTGAAGAACTGAAAAAATA<br>TCGTCCTGACATTTTAGACGCGTGTGAAAAGGCAATGAGTGCAGTGGACCCA<br>GACTTAAACTTTATTCGTGTAGACGAAGAGGCTTTCTTGGCATGTCCAGAAGA<br>ATCCGTTGACTATGCCGTGATGGAGCGTACGGCTGATGCGGTAGTTGTCCCA<br>ATGGACGCTGGATGGTCCGATGTTGGCAGCTGGTCATCGCTTTGGGAAATTA<br>GCGCCCACACCGCGGAAGGAAATGTATGTCACGGCGACGTGATTAACCATAA<br>AACAGAAAATTCATACGTTTATGCGGAATCCGGCTTGGTCACTACTGTCGGAG<br>TGAAGGATTTGGTTGTGGTGCAAACGAAAGATGCAGTATTAATTGCGGACCG<br>CAATGCTGTCCAAGATGTGAAGAAAGTAGTTGAACAGATTAAGGCTGATGGT<br>CGTCACGAGCATCGCGTCCACCGCGAAGTATATCGCCCATGGGGAAAGTAT<br>GACTCTATTGATGCGGGTGACCGCTATCAAGTTAAACGTATCACAGTCAAACC<br>CGGTGAGGGGCTTTCGGTGCAAATGCATCATCATCGCGCAGAGCATTGGGTA<br>GTGGTTGCGGGTACTGCGAAAGTAACAATTGATGGCGATATCAAGTTGCTTG<br>GCGAAAATGAGTCAATTTACATCCCGCTGGGCGCGACACACTGTCTTGAGAA<br>TCCGGGGAAAATCCCATTGGATTTAATCGAAGTGCGTTCTGGATCTTACTTAG<br>AGGAGGATGACGTTGTTCGTTTTGCTGATCGTTACGGACGCGTTtAAGTGAGT<br>AAGATAGGAT |

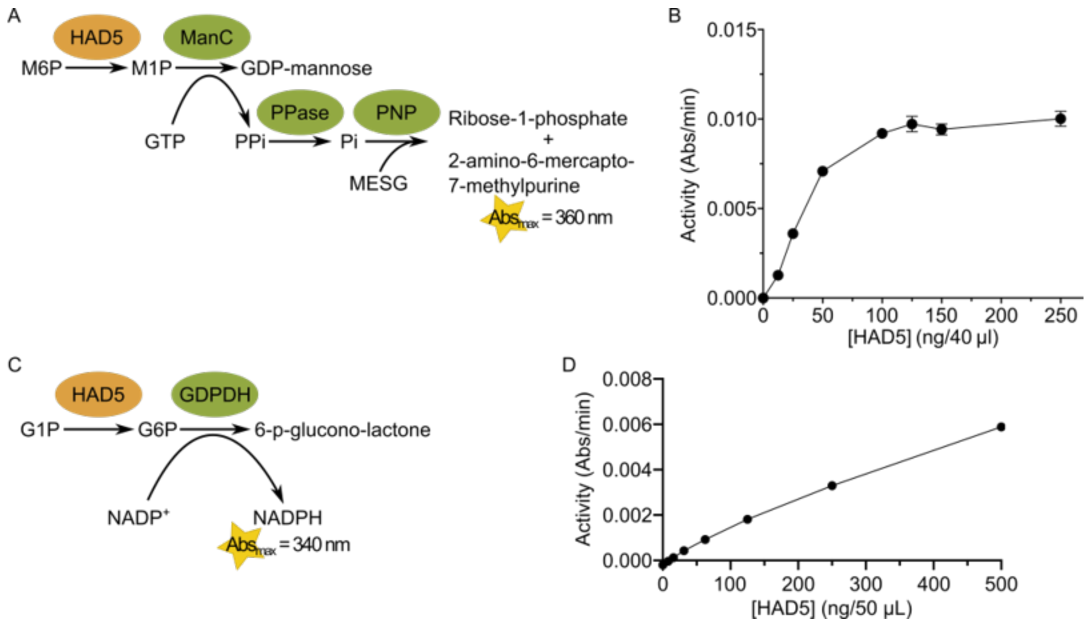


**Figure S1. Enzymatic activity assays**. **(A)** Schematic of the phosphomannomutase (PMM) activity assay, in which HAD5 converts mannose 6-phosphate (M6P) to mannose 1-phosphate (M1P). A series of linked-enzyme steps translates that activity into a spectrophotometric signal at 360 nm. **(B)** Graph depicting enzymatic activity in the PMM assay with varying HAD5 concentration, demonstrating that our chosen value of 50 ng / 40 µL reaction is within the linear range of the assay with respect to enzyme concentration. **(C)** Schematic of the phosphoglucomutase (PGM) activity assay, in which HAD5 converts glucose 1-phosphate (G1P) to glucose 6-phosphate (G6P), which is then used by glucose 6-phosphate dehydrogenase (G6PDH) to generate NADPH, which can be measured at 340 nm. **(D)** Graph depicting enzymatic activity in the PGM assay with varying HAD5 concentration, demonstrating that our chosen value of 50 ng / 50 µL reaction is within the linear range of the assay with respect to enzyme concentration.

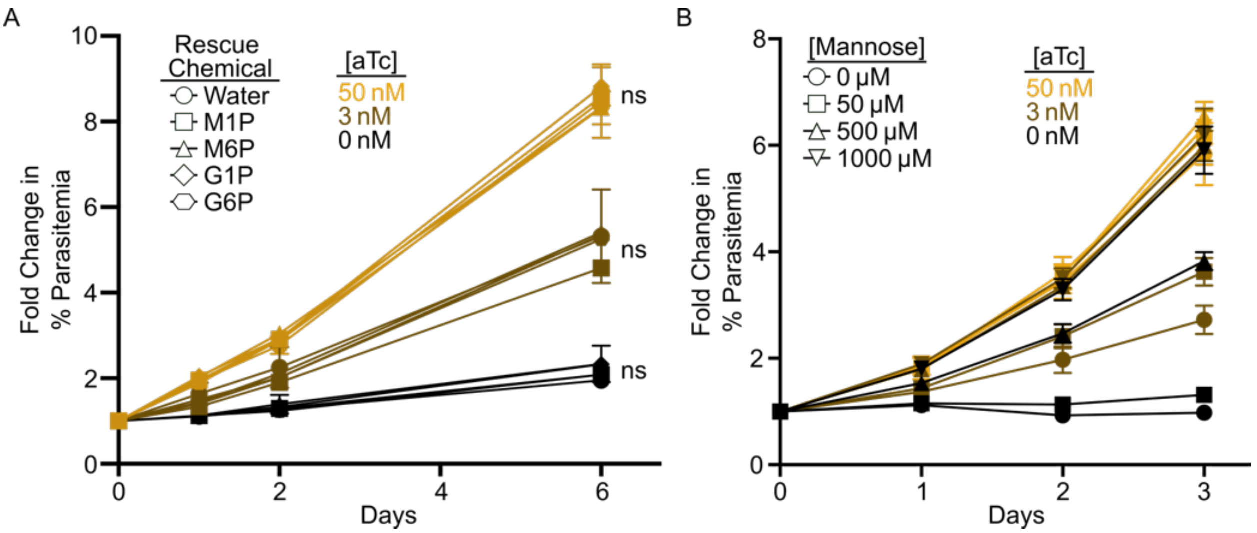


**Figure S2. Chemical rescue of parasite growth. (A)** HAD5$^{KD}$ parasite growth was measured in the presence of varying anhydrotetracycline (aTc) concentrations with chemical rescue by four different chemicals: mannose 1-phosphate (M1P), mannose 6-phosphate (M6P), glucose 1-phosphate (G1P), and glucose 6-phosphate (G6P) at 20 μM concentration. Data depicts the mean ± standard error of the mean (SEM) of duplicate experiments and represents fold change in parasitemia over time. Statistics were performed on the Day 6 parasitemia, using a two-way ANOVA with Dunnett's multiple comparisons test, with individual variances computed for each comparison. In all cases, chemical rescues were compared to the vehicle control within a given aTc concentration, with no significant rescue of growth observed (ns = not significant). **(B)** Shown are the complete data of the fold change in parasitemia over time when HAD5$^{KD}$ parasites are grown in varying aTc and D-mannose concentrations. Data represent mean ± SEM of three independent experiments with technical duplicates. The data from the Day 3 time point was used to generate Figure 2B.

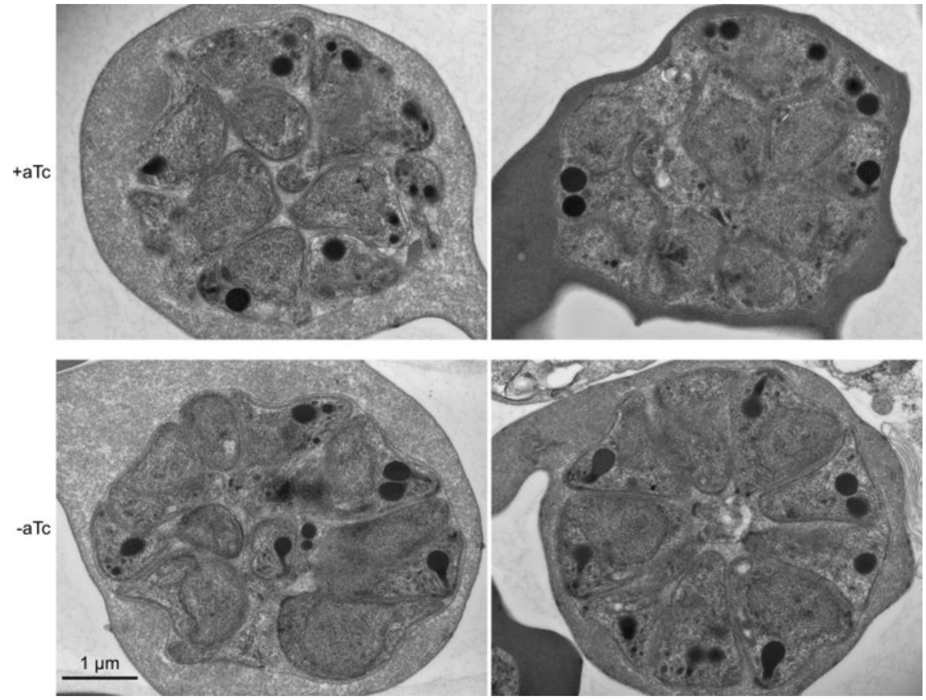


**Figure S3. Ultrastructures of HAD5$^{KD}$ schizonts by transmission electron micrograph.** Transmission electron microscopy of highly synchronized schizont parasites showing successful schizogony in parasites grown under -aTc conditions compared to +aTc.

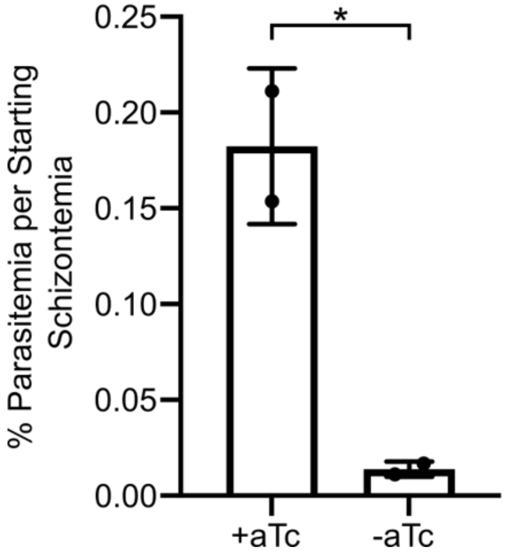


**Figure S4. HAD5 KD Parasites are deficient in reinvasion**. Schizont-stage parasites 40-44 hours post invasion, grown in ±aTc conditions, were E64-arrested as late schizonts for 8 hours, mechanically lysed, and allowed to reinvade over fresh red blood cells. 24 hours later, parasitemia was assessed by flow cytometry and normalized to pre-lysis schizontemia, demonstrating a deficiency in reinvasion by parasites grown in –aTc conditions. Statistics were performed using an unpaired two-tailed t-test. $^*p=0.028$

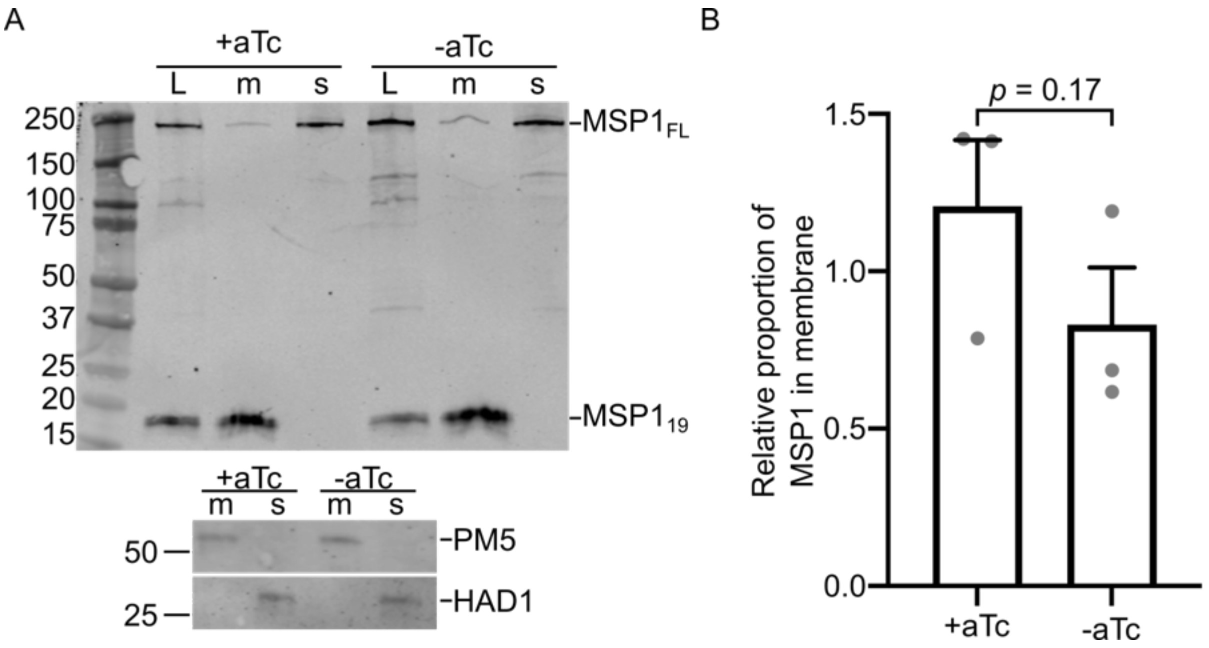


**Figure S5. TritonX-114 partition of MSP1**. **(A)** Representative Western blot of whole lysate (L), membranous fraction (m), and soluble fraction (s) of 40-44hr parasites grown in ±aTc, and blotted for MSP1. Full Length (FL) and 19 kDa fragments of MSP1 are indicated. Plasmepsin 5 (PM5) and HAD1 were used as membrane and soluble controls, respectively. Molecular Weight ladder sizes are indicated in kDa on the left. **(B)** Quantification of A. Full Length and $MSP1_{19}$ were summed in each lane, and the relative signal in membranous lanes were compared to that of whole lysate. Data represent mean + SEM of three independent experiments. Statistics were performed using a paired, two-tailed t-test.

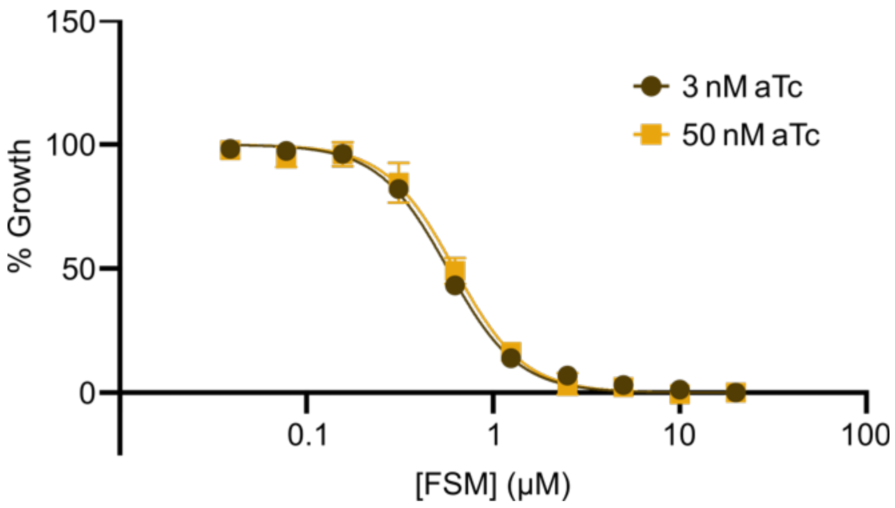


**Figure S6. Knockdown of HAD5 has no effect on FSM sensitivity**. 72-hour dose-response curves of asynchronous parasites treated with fosmidomycin in either intermediate or saturating aTc concentrations. % Growth was relative to a vehicle control. The $EC_{50}$ ± SEM (in μM) for each condition were: 3 nM, 0.57 ± 0.02; 50 nM, 0.62 ± 0.04. Not significant ($p$ = 0.39) by unpaired t-test.

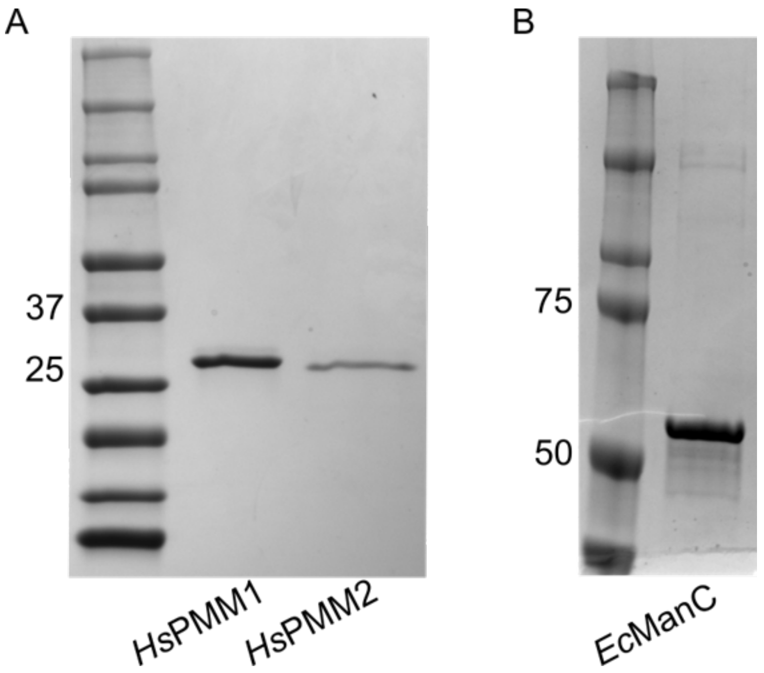


**Figure S7. Purified recombinant proteins**. Coomassie-stained SDS-PAGE gels of final purified forms of recombinant *Hs*PMM1, *Hs*PMM2 **(A)** and *Ec*ManC **(B)**.

A

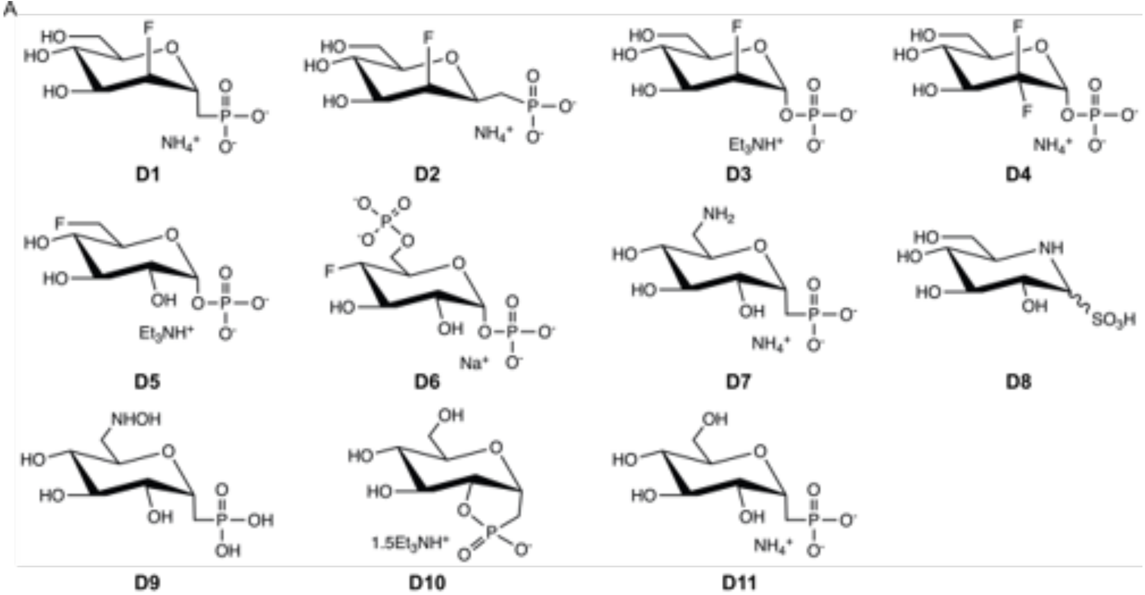
D1
D2
D3
D4
D5
D6
D7
D8
D9
D10
D11

B

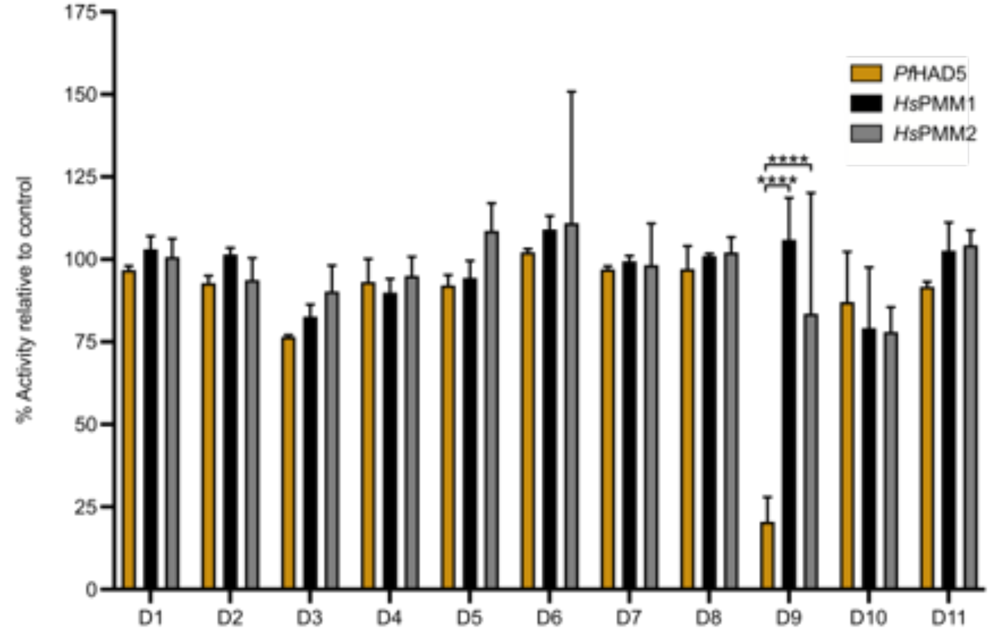
% Activity relative to control
PfHAD5
HsPMM1
HsPMM2
D1 D2 D3 D4 D5 D6 D7 D8 D9 D10 D11

C

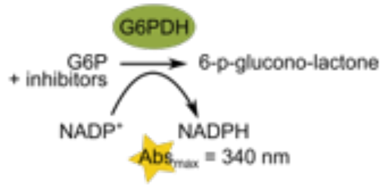
G6PDH
G6P
+ inhibitors
6-p-glucono-lactone
NADP+
NADPH
Abs max = 340 nm

D

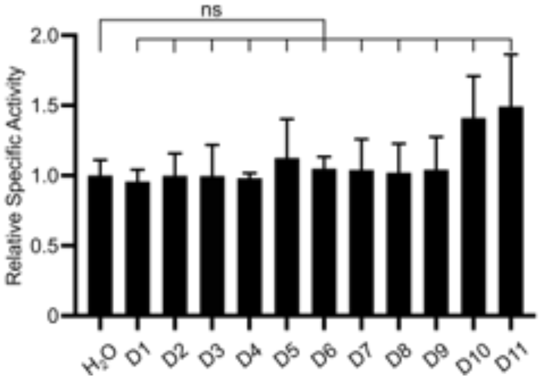
ns
Relative Specific Activity
H2O D1 D2 D3 D4 D5 D6 D7 D8 D9 D10 D11

**Figure S8. Structures and evaluation of compounds D1-D11. (A)** Structures of the 11 compounds tested for inhibition against *Pf*HAD5, *Hs*PMM1, and *Hs*PMM2. **(B)** Activity of recombinant *Pf*HAD5, *Hs*PMM1, and *Hs*PMM2 when treated with 125 μM of the indicated compounds as a percentage of treatment with a vehicle control. Data represent the mean ± SEM of 3 independent experiments with technical replicates. Statistics were performed with a two-way ANOVA using Tukey's test for multiple comparisons. ****$p<0.0001$. All other comparisons between enzymes for a given compound were not significant. **(C)** Diagram of the amended assay to assess compound inhibition of downstream components of the assay. Assays were commenced with the addition of substrate (G6P). **(D)** Quantification of assay shown in C. Assay activity is depicted relative to treatment with a vehicle control. Data represent the mean ± SEM of 3 independent experiments with technical replicates. Statistics were performed with an ordinary one-way ANOVA using Dunnett's test for multiple comparisons with single pooled variance. ns = not significant.

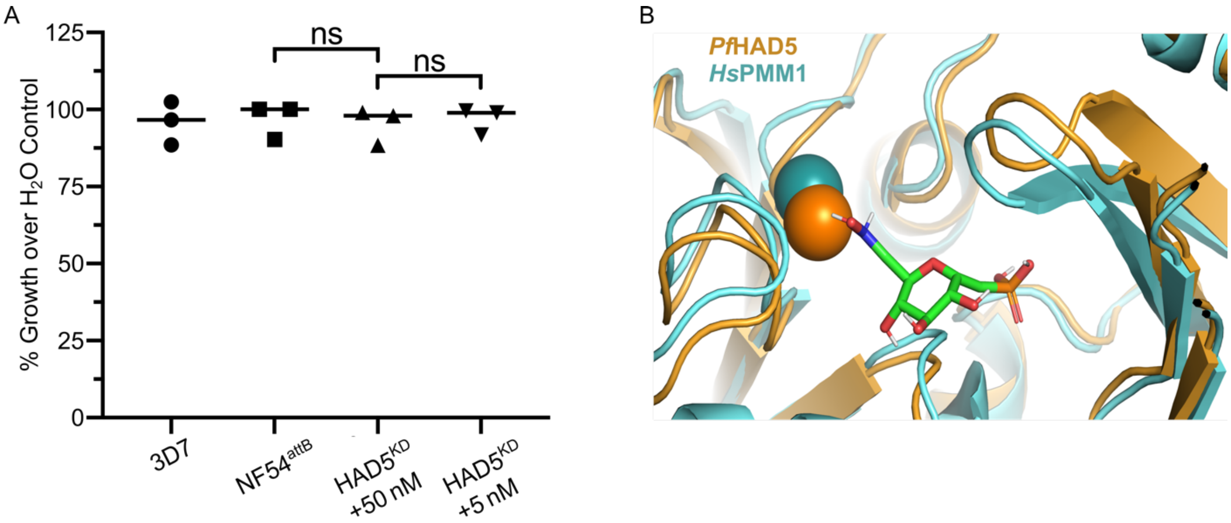


**Figure S9. Compound D9 does not inhibit parasites in culture. (A)** Displayed is the percent growth after 72 hours of parasites (starting parasitemia ~1.2%) when treated with 100 µM compound D9, compared to vehicle control. The experiment was performed on wild type parasites of two different strains (3D7 and NF54), as well as HAD5$^{KD}$ parasites ±aTc. Three independent experiments, each with technical replicates, were performed. Statistics were performed with an ordinary one-way ANOVA using Tukey's test for multiple comparisons, ns = not significant. **(B)** Shown is the D9 compound (green), computationally docked to *Pf*HAD5 crystal structure (orange), highlighting its location in the binding pocket, compared to subtle differences in the binding pocket of *Hs*PMM1 (cyan). Mg2+ ions are depicted from *Pf*HAD5 (dark orange) and *Hs*PMM1 (dark cyan) as spheres.

### Synthesis of phosphonate analogue of 6NHOH-G1CP (D9)

**Scheme 1. Synthesis of phosphonate analogue of 6NHOH-G1CP (D9)**

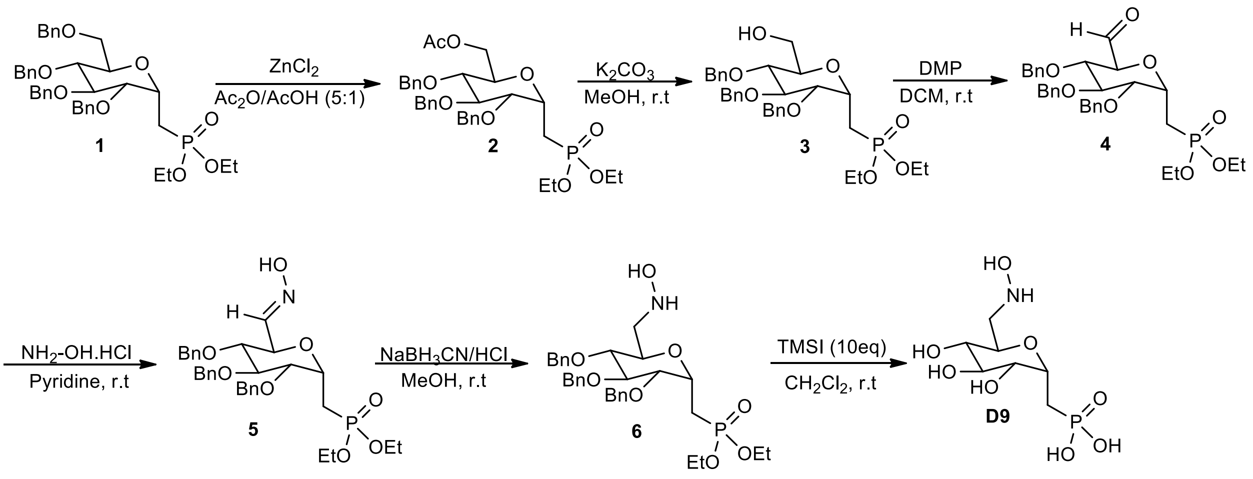


The synthesis of α-D-glucose-1-phosphonate **D9** analogue commenced with the selective debenzylation-acetolysis (105) of the benzyl C-6 substituent of diethyl-C-(1-Deoxy 2,3,5,6-tetra-O-benzyl-α-D-gluctopyranosyl) methanephosphonate (**1**) using a solution of freshly fused $ZnCl_2$ (5.2 equiv) in 1:5 HOAc–Ac2O in 98% yield. (Scheme 1). We observed that it was important to keep the temperature at 0 °C during the course of the reaction (30 min). The precursor phosphonate of **1** was synthesized according to our previously reported in three steps starting from 2,3,4,6-tetra-*O*-benzyl-D-glucopyranose (106). Conventional de-O-acetylation of **2** with potassium carbonate in methanol gave the alcohol **3** in high yield. Although deacetylation of the acetyl protecting group at C-6 in **2** was first attempted using a mixture 1:3:7 TEA-$H_2O$-MeOH, but only poor yields of alcohol **2** was obtained mainly due to low solubility of **2**. Synthetic alcohol **3** was converted to the corresponding aldehyde **4** by using the Dess-Martin periodinane (DMP) in 98% yield (107). First we examined reaction with using 1 equivalent of DMP, but the yield was low, then we increased amounts of DMP to 1.5 and 2 equivalents. Our optimal result was obtained when we used 1.5 equivalents of DMP. Without further purification, condensation of aldehyde **4** with hydroxylamine hydrochloride in pyridine afforded the desired oxime **5** in 94% yield (107). Further reduction of **5** with sodium cyanoborohydride ($NaBH_3CN$) under acidic condition provided the hydroxylamine **6** in 78% yield (107). Deprotection of both benzyl ether and ethyl ester groups was readily accomplished with an excess of iodotrimethylsilane (TMSI) (106, 108). For preparation of target phosphonate **D9**, different quantities of TMSI were examined. With 10 equivalents of TMSI compound **6,** was converted to the final product **D9** in 95% yield. It should be mentioned, that when more than 10 equivalents of TMSI was used, the reaction led to decomposition.

**((2R,3R,4S,5R,6S)-3,4,5-tris(benzyloxy)-6-((diethoxyphosphoryl)methyl)tetrahydro-2H-pyran-2-yl)methyl acetate (2).**

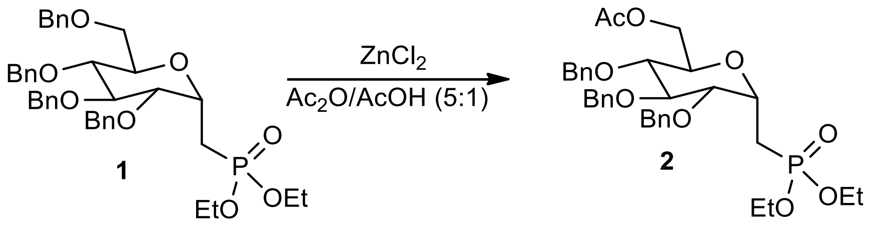


A solution of $ZnCl_2$ (708 mg, 5.2 eq, 5.2 mmol) in $Ac_2O$/AcOH (5:1, 12 mL) was cooled to 0 °C then a solution of **1** (674 mg, 1mmol) in $Ac_2O$/AcOH (5:1, 12 mL) was added dropwise. The solution was stirred for overnight at room temperature under $N_2$. After evaporation, the residue was suspended in water (20 mL) and extracted with DCM (40 mL x 2). The combined organic extracts were dried ($MgSO_4$), filtered and concentrated. Purification by flash chromatography (Hexanes-EtOAc, 1:1) afforded compound **2** (660 mg, 98%) as a colorless liquid; Rf 0.50 (Hexanes-EtOAc, 1:3). $^1$H NMR (500 MHz, $CDCl_3$), $\delta_H$ (ppm) 1.35 (6H, td, $^3J_{HH}$ = 7.1 Hz, $^4J_{HP}$ = 2.4 Hz, $2OCH_2CH_3$), 2.06 (3H, s, $CH_3CO$), 2.20-2.26 (2H, m, $CH_2$-P), 3.57 (1H, t, $^3J_{HH}$ = 9.1 Hz, H-4), 3.72-3.75 (2H, m, H-2, H-3), 3.80 (1H, dt, $^3J_{HH}$ = 9.7 Hz, $^3J_{HH}$ = 2.8 Hz, H-5), 4.10-4.16 (4H, m, $2OCH_2CH_3$), 4.24 (1H, dd, $^2J_{HH}$ = 12.0 Hz, $^3J_{HH}$ = 2.1 Hz, H-6a or H-6b), 4.45 (1H, dd, $^2J_{HH}$ = 12.0 Hz, $^3J_{HH}$ = 3.6 Hz, H-6a or H-6b), 4.35-4.56 (1H, m, H-1), 4.58 (1H, d, $^2J_{HH}$ = 10.9 Hz, $CH_2$ of Bn), 4.68 (1H, d, $^2J_{HH}$ = 11.5 Hz, $CH_2$ of Bn), 4.71 (1H, d, $^2J_{HH}$ = 11.5 Hz, $CH_2$ of Bn), 4.81 (1H, d, $^2J_{HH}$ = 11.5 Hz, $CH_2$ of Bn), 4.87 (1H, d, $^2J_{HH}$ = 10.9 Hz, $CH_2$ of Bn), 4.94 (1H, d, $^2J_{HH}$ = 10.9 Hz, $CH_2$ of Bn), 7-28-7.36 (15H, m, 3Ph). $^{13}$C NMR (125 MHz, $CDCl_3$), $\delta_C$ (ppm) 16.69 (d, $^3J_{CP}$ = 3.7 Hz, $2OCH_2CH_3$), 16.73 (d, $^3J_{CP}$ = 3.7 Hz, $2OCH_2CH_3$), 21.08 ($CH_3CO$), 22.87 (d, $^1J_{CP}$ = 144.0 Hz, $CH_2$-P), 61.81 (d, $^2J_{CP}$ = 6.4 Hz, $OCH_2CH_3$), 62.03 (d, $^2J_{CP}$ = 6.1 Hz, $OCH_2CH_3$), 63.31 ($CH_2$-6), 69.88 (d, $^2J_{CP}$ = 5.1 Hz, C-1), 70.49 (C-5), 73.34, 75.14, 75.58 (3$CH_2$ of Bn), 77.40 (C-4), 79.31 (d, $^3J_{CP}$ = 12.7 Hz, C-2), 77.40 (C-3), 127.98, 128.14, 128.29, 128.67, 128.71, 137.95, 137.99, 138.53 (3Ph), 170.92 (C=O).$^{31}$P NMR (121 MHz, $CDCl_3$), $\delta_P$ (ppm) 28.88. HRMS (ESI, Positive Mode) calcd for $C_{34}H_{43}NaO_9P$ [M + Na]$^+$ 649.2537, found 649.2534.

**((2R,3R,4S,5R,6S)-3,4,5-tris(benzyloxy)-6-((diethoxyphosphoryl)methyl)tetrahydro-2H-pyran-2-yl)methyl acetate (3).**

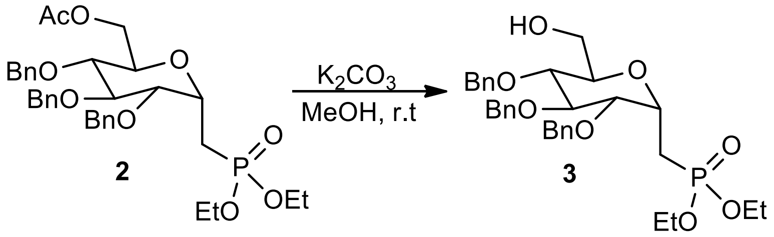


To a solution of **2** (626 mg, 1 mmol) in MeOH (30 mL), $K_2CO_3$ (500mg, 3.6 mmol) was added at room temperature. The resulting solution was then stirred overnight. After evaporation, the residue was suspended in water (30 mL) and extracted with DCM (40 mL x 2). The combined organic extracts were dried ($MgSO_4$), filtered and concentrated to provide the crude product (543 mg, 93%) as a colorless liquid which was used in subsequent reactions without further purification (Rf 0.22 (Hexanes-EtOAc, 1:3). $^1$H NMR (500 MHz, $CDCl_3$), $\delta_H$ (ppm) 1.35 (6H, t, $^3J_{HH}$ = 7.1 Hz, $2OCH_2CH_3$), 1.78 (1H, brs, OH), 2.04-2.12 (1H, m, $CH_2$-P), 2.32-2.41 (1H, m, $CH_2$-P), 3.40 (1H, t, $^3J_{HH}$ = 8.1 Hz, H-4), 3.60-3.67 (1H, m, H-2), 3.73-3.81 (3H, m, H-3, H-6a, H-6b), 3.85 (1H, td, $^3J_{HH}$ = 8.3 Hz, $^3J_{HH}$ = 2.6 Hz, H-5), 4.12-4.17 (4H, m, $2OCH_2CH_3$), 4.43-4.45 (1H, m, H-1), 4.61-4.83 (6H, m, $3CH_2$ of Bn), 7-29-7.40 (15H, m, 3Ph). $^{13}$C NMR (125 MHz, $CDCl_3$), $\delta_C$ (ppm) 16.62 (d, $^3J_{CP}$ = 5.8 Hz, $2OCH_2CH_3$), 24.38 (d, $^1J_{CP}$ = 144.1 Hz, $CH_2$-P), 61.97 (d, $^2J_{CP}$ = 5.9 Hz, $OCH_2CH_3$), 62.00 ($CH_2$-6), 62.25 (d, $^2J_{CP}$ = 6.4 Hz, $OCH_2CH_3$), 68.66 (d, $^2J_{CP}$ = 5.6 Hz, C-1), 73.42 ($CH_2$ of Bn), 73.89 (C-5), 74.76, 75.14 ($2CH_2$ of Bn), 77.42 (C-4), 78.88 (d, $^3J_{CP}$ = 13.7 Hz, C-2), 80.79 (C-3), 127.95, 128.02, 128.09, 128.17, 128.19, 128.64, 128.67, 128.70, 137.98, 138.11, 138.47 (3Ph).$^{31}$P NMR (121 MHz, $CDCl_3$), $\delta_P$ (ppm) 29.79. HRMS (ESI, Positive Mode) calcd for $C_{32}H_{41}NaO_8P$ [M + Na]$^+$ 607.2431, found 607.2420.

**Diethyl(((2S,3R,4S,5S,6S)-3,4,5-tris(benzyloxy)-6-formyltetrahydro-2H-pyran-2-yl) methyl) phosphonate (4).**

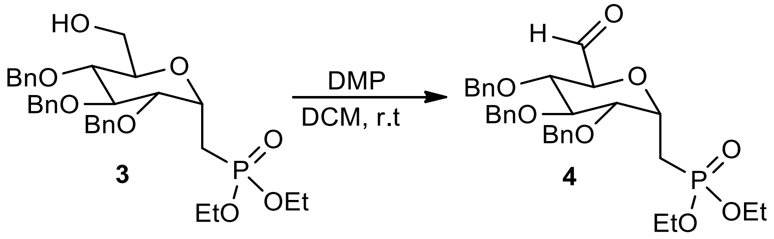


To a solution of **3** (584 mg, 1 mmol) in anhydrous $CH_2Cl_2$ (30 mL), Dess-Martin periodinane (636 mg, 1.5 mmol, 1.5 equiv) was added at room temperature. The resulting solution was then stirred at same temperature for 4h under nitrogen. The solution was diluted $CH_2Cl_2$ (30 mL), and then saturated $NaHCO_3$(aq) and saturated $Na_2S_2O_3$(aq) were added to the reaction sequentially. The resulting mixture was stirred for another 30 min at room temperature. The organic layer was separated, and the aqueous layer was extracted with EtOAc. The combined organic extracts were dried ($MgSO_4$), filtered and concentrated to provide the crude product (547 mg, 94%) as a colorless liquid which was used in subsequent reactions without further purification (Rf 0.30 (Hexanes-EtOAc, 1:3). $^1$H NMR (500 MHz, $CDCl_3$), $\delta_H$ (ppm) 1.28-1.35 (6H, m, 2OCH$_2$C$H_3$), 2.11-2.19 (1H, m, $CH_2$-P), 2.31-2.39 (1H, m, $CH_2$-P), 3.52 (1H, dd, $^3J_{HH}$ = 5.3 Hz, $^4J_{HP}$ = 3.3 Hz, H-2), 3.76 (1H, t, $^3J_{HH}$ = 5.3 Hz, H-3), 3.81 (1H, t, $^3J_{HH}$ = 4.8 Hz, H-4), 4.09-4.18 (4H, m, 2OC$H_2$CH$_3$), 4.28 (1H, d, $^3J_{HH}$ = 4.7 Hz, H-5), 4.46 (1H, d , $^2J_{HH}$ = 11.6 Hz, $CH_2$ of Bn), 4.53-4.57 (3H, m, H-1 and $CH_2$ of Bn ), 4.62-4.70 (3H, m, $CH_2$ of Bn ), 7-20-7.36 (15H, m, 3Ph), 9.83 (1H, s, CHO). $^{31}$P NMR (121 MHz, $CDCl_3$), $\delta_P$ (ppm) 28.67.

**Diethyl (((2S,3R,4S,5R,6R)-3,4,5-tris(benzyloxy)-6-((E)-(hydroxyimino)methyl)tetrahydro-2H-pyran-2-yl) methyl)phosphonate (5).**

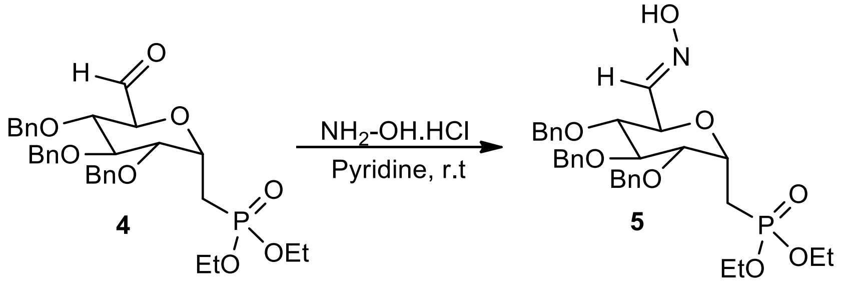


To a solution of **4** (582 mg, 1 mmol) in anhydrous pyridine (15 mL), hydroxylamine hydrochloride (104 mg, 1.5 mmol, 1.5 equiv) was added at room temperature. The resulting solution was then stirred at same temperature for 5h and then pyridine was removed at 50 °C under high vacuum. Water (30 mL) was added to the residue and the mixture was extracted with $CH_2Cl_2$ (30 mL x 2). The combined organic extracts were dried ($MgSO_4$), filtered and concentrated. Purification by flash column chromatography (Hexanes-EtOAc, 1:2) afforded compound **5** as a mixture of E/Z isomers (561 mg, 94%) as a colorless liquid; Rf 0.41 (Hexanes-EtOAc, 1:3). $^{1}$H NMR (500 MHz, $CDCl_3$), $\delta_H$ (ppm) 1.29-1.37 (6H, m, 2OCH$_2$C*H*$_3$), 2.07-2.50 (2H, m, CH$_2$-P), 3.43-3.80 (3H, m, H-2, H-3, H-4), 4.06-4.19 (4H, m, 2OC*H*$_2$CH$_3$), 4.23-4.36 (1H, m, H-5), 3.48-4.55 (1H, m, H-1), 4.57-4.95 (6H, m, 3CH$_2$ of Bn), 7-15-7.37 (16H, m, 3Ph, CH=N), 9.38 (1H, brs, OH). $^{13}$C NMR (125 MHz, $CDCl_3$), $\delta_C$ (ppm) 16.52 (d, $^{3}J_{CP}$ = 6.5 Hz, OCH$_2$*C*H$_3$), 16.60 (d, $^{3}J_{CP}$ = 6.4 Hz, OCH$_2$*C*H$_3$), 22.36 (d, $^{1}J_{CP}$ = 143.2 Hz, CH$_2$-P), 62.05 (d, $^{2}J_{CP}$ = 6.4 Hz, O*C*H$_2$CH$_3$), 62.20 (d, $^{2}J_{CP}$ = 5.7 Hz, O*C*H$_2$CH$_3$), 70.10 (d, $^{2}J_{CP}$ = 5.4 Hz, C-1), 71.31 (C-5), 73.39, 75.24, 75.43 (3CH$_2$ of Bn), 78.81 (d, $^{3}J_{CP}$ = 12.9 Hz, C-2), 80.13, 80.71 (C-3, C-4), 127.74, 127.82, 128.05, 128.12, 128.21, 128.51, 128.63, 128.70, 137.91, 138.01, 138.71 (3Ph), 148.14 (CH=N).$^{31}$P NMR (121 MHz, $CDCl_3$), $\delta_P$ (ppm) 28.96 (minor isomer), 30.46 (major isomer). HRMS (ESI, Positive Mode) calcd for $C_{32}H_{40}NNaO_8P$ [M + Na]$^{+}$ 620.2384, found 620.2391.

**Diethyl (((2S,3R,4S,5R,6R)-3,4,5-tris(benzyloxy)-6-((hydroxyamino)methyl)tetrahydro-2H-pyran-2-yl)methyl)phosphonate (6).**

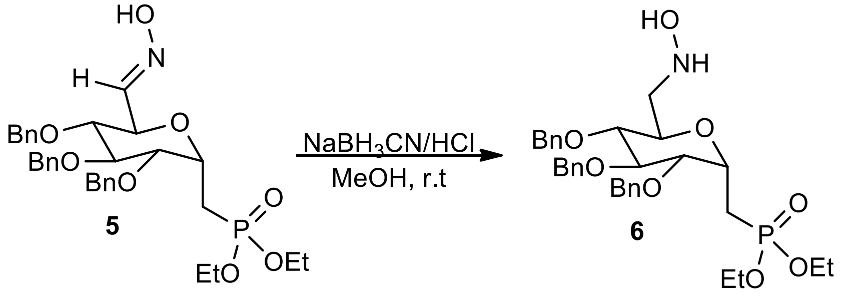


To a solution of **5** (597 mg, 1 mmol) and $NaBH_3CN$ (125 mg, 2 mmol, 2 equiv) in MeOH (30 mL), a solution of HCl (6N in MeOH) was added dropwise at 0 °C until pH 1-3. The resulting solution was then stirred at room temperature for 10h. The reaction was quenched by the addition of saturated $NaHCO_3$ (aq) and the mixture was extracted with EtOAc (40 mL x 2). The combined organic extracts were dried ($MgSO_4$), filtered and concentrated. Purification by flash column chromatography ($CH_2Cl_2$-MeOH, 40:1) afforded compound **5** (467 mg, 78%) as a colorless liquid; Rf 0.50 ($CH_2Cl_2$-MeOH, 40:1). $^1$H NMR (500 MHz, $CDCl_3$), $\delta_H$ (ppm) 1.35 (6H, td, $^3J_{HH}$ = 7.1 Hz, $^4J_{HP}$ = 1.7 Hz, 2OCH$_2$C$H_3$), 2.13-2.21 (1H, m, $CH_2$-P), 2.33-2.42 (1H, m, $CH_2$-P), 2.82 (1H, dd, $^2J_{HH}$ = 13.6 Hz, $^3J_{HH}$ = 9.5 Hz, H-6a or H-6b), 3.27 (1H, t, $^3J_{HH}$ = 9.3 Hz, H-4), 3.40 (1H, dd, $^2J_{HH}$ = 13.6 Hz, $^3J_{HH}$ = 1.7 Hz, H-6a or H-6b), 3.70-3.79 (2H, m, H-2, H-3), 4.00 (1H, dd, $^3J_{HH}$ = 9.5 Hz, $^3J_{HH}$ = 1.7 Hz, H-5), 4.09-4.22 (4H, m, 2OC$H_2$CH$_3$), 4.35-4.41 (1H, m, H-1), 4.62 (1H, d, $^2J_{HH}$ = 11.0 Hz, $CH_2$ of Bn), 4.64 (1H, d, $^2J_{HH}$ = 11.7 Hz, $CH_2$ of Bn), 4.76 (1H, d, $^2J_{HH}$ = 11.7 Hz, $CH_2$ of Bn), 4.83 (1H, d, $^2J_{HH}$ = 10.9 Hz, $CH_2$ of Bn), 4.87 (1H, d, $^2J_{HH}$ = 11.0 Hz, $CH_2$ of Bn), 4.92 (1H, d, $^2J_{HH}$ = 10.9 Hz, $CH_2$ of Bn), 7-29-7.36 (15H, m, 3Ph). $^{13}$C NMR (125 MHz, $CDCl_3$), $\delta_C$ (ppm) 16.61 (d, $^3J_{CP}$ = 4.8 Hz, 2OCH$_2$C$H_3$), 16.64 (d, $^3J_{CP}$ = 4.6 Hz, 2OCH$_2$C$H_3$), 24.16 (d, $^1J_{CP}$ = 145.7 Hz, $CH_2$-P), 55.59 ($CH_2$-6), 61.96 (d, $^2J_{CP}$ = 6.3 Hz, OC$H_2$CH$_3$), 62.68 (d, $^2J_{CP}$ = 6.5 Hz, OC$H_2$CH$_3$),68.97 (C-5), 68.93 (d, $^2J_{CP}$ = 5.6 Hz, C-1), 73.70, 75.20, 75.71 (3$CH_2$ of Bn), 79.63 (d, $^3J_{CP}$ = 13.4 Hz, C-2), 80.80 (C-4), 82.13 (C-3) 172.89, 128.00, 128.04, 128.07, 128.09, 128.17, 128.60, 128.71, 138.00, 138.07, 138.58 (3Ph).$^{31}$P NMR (121 MHz, $CDCl_3$), $\delta_P$ (ppm) 28.83. HRMS (ESI, Positive Mode) calcd for $C_{32}H_{43}NO_8P$ [M]$^+$ 600.2721, found 600.2700.

**(((2S,3R,4S,5S,6R)-3,4,5-Trihydroxy-6-((hydroxyamino)methyl)tetrahydro-2H-pyran-2-yl) methyl)phosphonic acid (D9).**

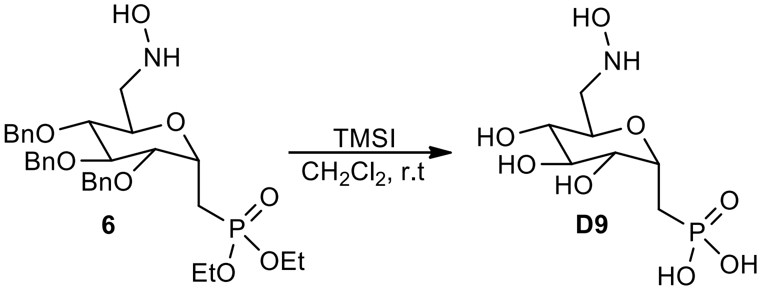


To a solution of **6** (180 mg, 0.3 mmol) in anhydrous $CH_2Cl_2$ (3 mL), iodotrimethylsilane (427 µL, 3.0 mmol, 10 equiv) was added dropwise at 0 °C under nitrogen. The resulting solution was then stirred at room temperature for 3h. The reaction was quenched by the addition of methanol. The mixture was concentrated, dissolved in $H_2O$ (10 mL) and washed with diethyl ether (20mL x 5). The aqueous layer was lyophilized to afford the target **D9** as a colorless foam (78 mg, 95%). $^1$H NMR (500 MHz, $D_2O$), $\delta_H$ (ppm) 2.01-2.06 (1H, m, $CH_2$-P), 2.21-2.27 (1H, m, $CH_2$-P), 3.31 (1H, t, $^3J_{HH}$ = 9.5 Hz, H-4), 3.57 (1H, dd, $^2J_{HH}$ = 13.6 Hz, $^3J_{HH}$ = 10.0 Hz, H-6a or H-6b), 3.57-3.62 (2H, m, H-3, H-6a or H-6b), 3.70-3.75 (1H, m, H-2), 3.92 (1H, dd, $^3J_{HH}$ = 9.8 Hz, $^3J_{HH}$ = 2.3 Hz, H-5), 4.39-4.41 (1H, m, H-1). $^{13}$C NMR (125 MHz, $D_2O$), $\delta_C$ (ppm) 23.29 (d, $^1J_{CP}$ = 136.9 Hz, $CH_2$-P), 52.50 ($CH_2$-6), 66.28 (C-5), 70.55 (d, $^3J_{CP}$ = 12.9 Hz, C-2), 71.73 (C-4), 72.26 (d, $^2J_{CP}$ = 5.8 Hz, C-1), 72.60 (C-3). $^{31}$P NMR (121 MHz, $D_2O$), $\delta_P$ (ppm) 24.50. HRMS (ESI, Positive Mode) calcd for $C_7H_{17}NO_8P$ $[M+1]^+$ 274.0686, found 274.0694.

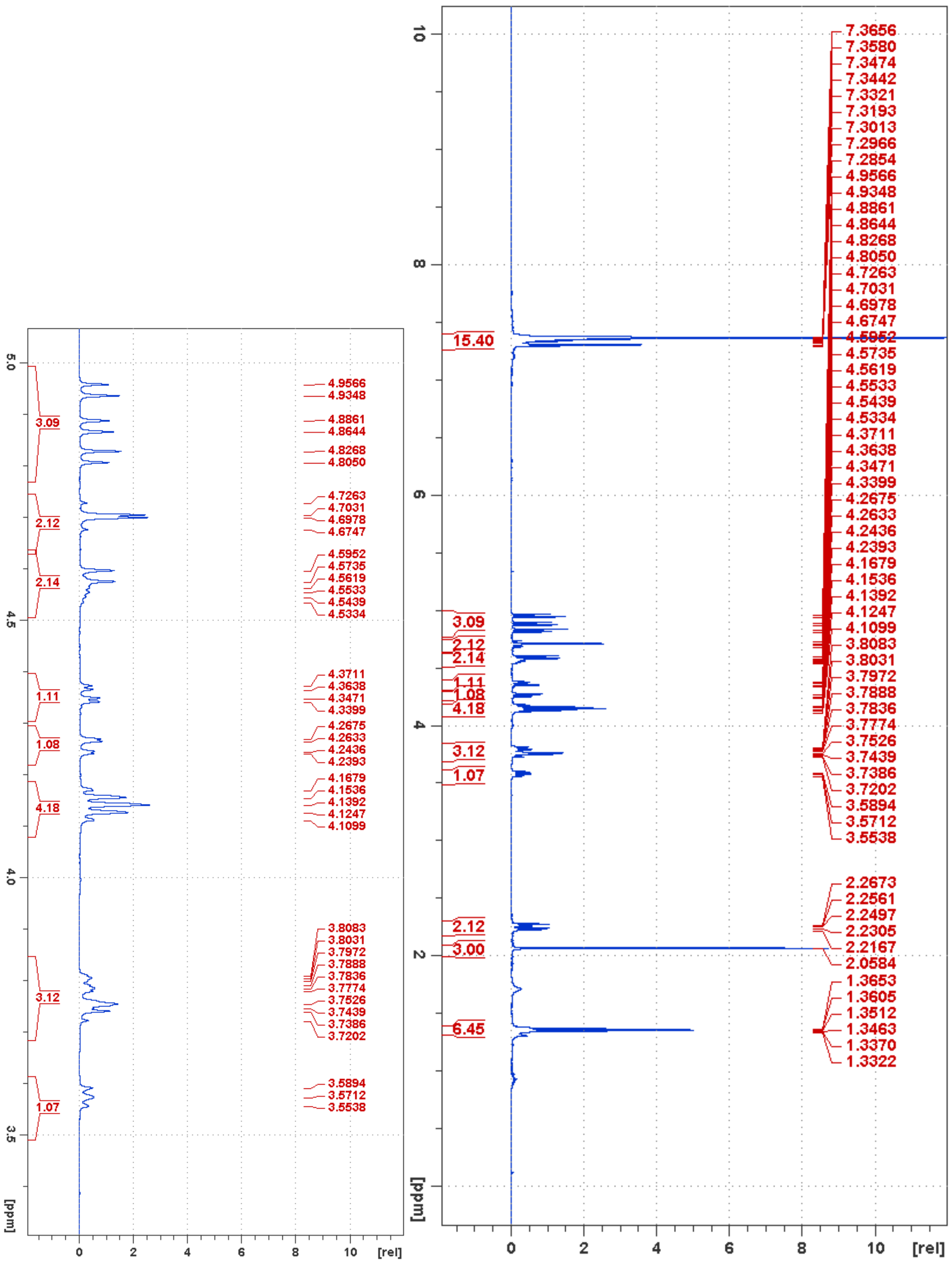


**Figure S10**. $^1$H NMR of compound **2**

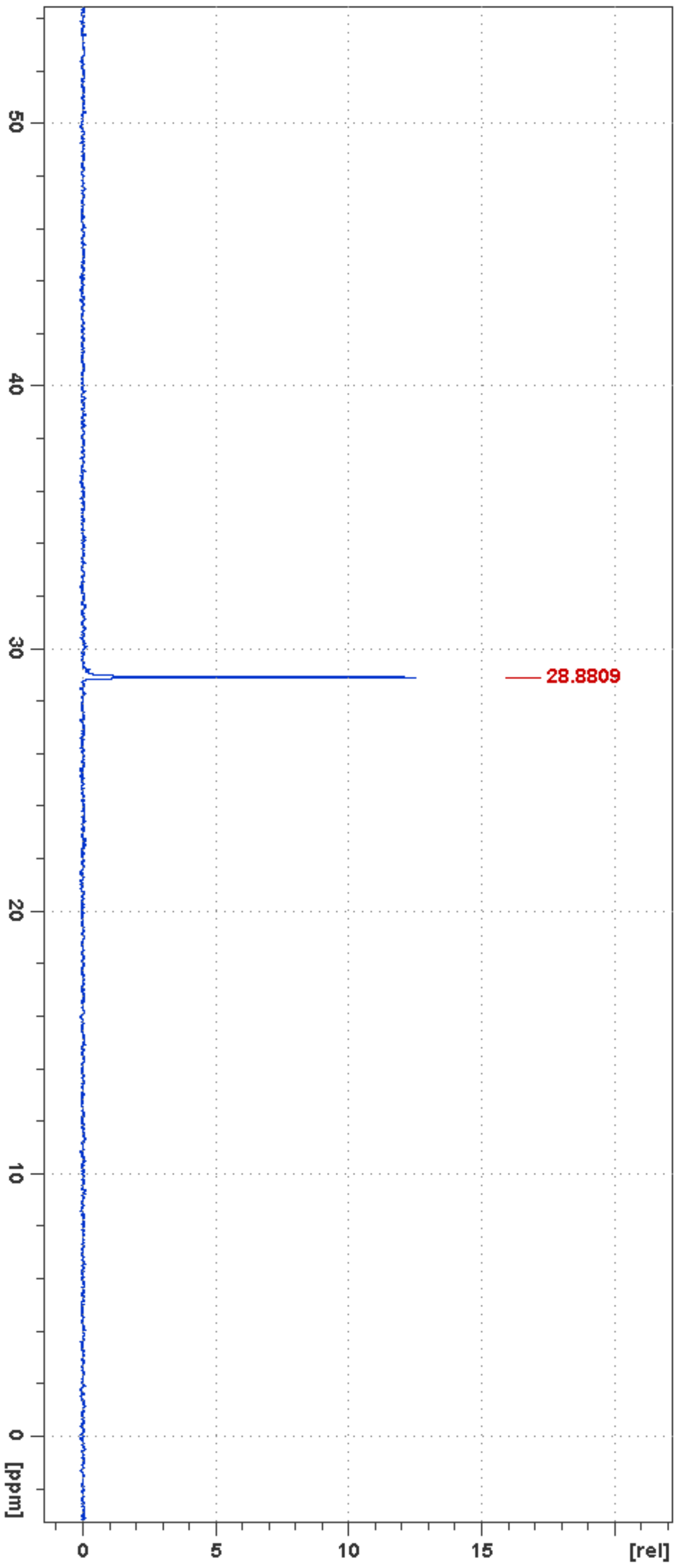


**Figure S11**. $^{31}$P NMR of compound **2**

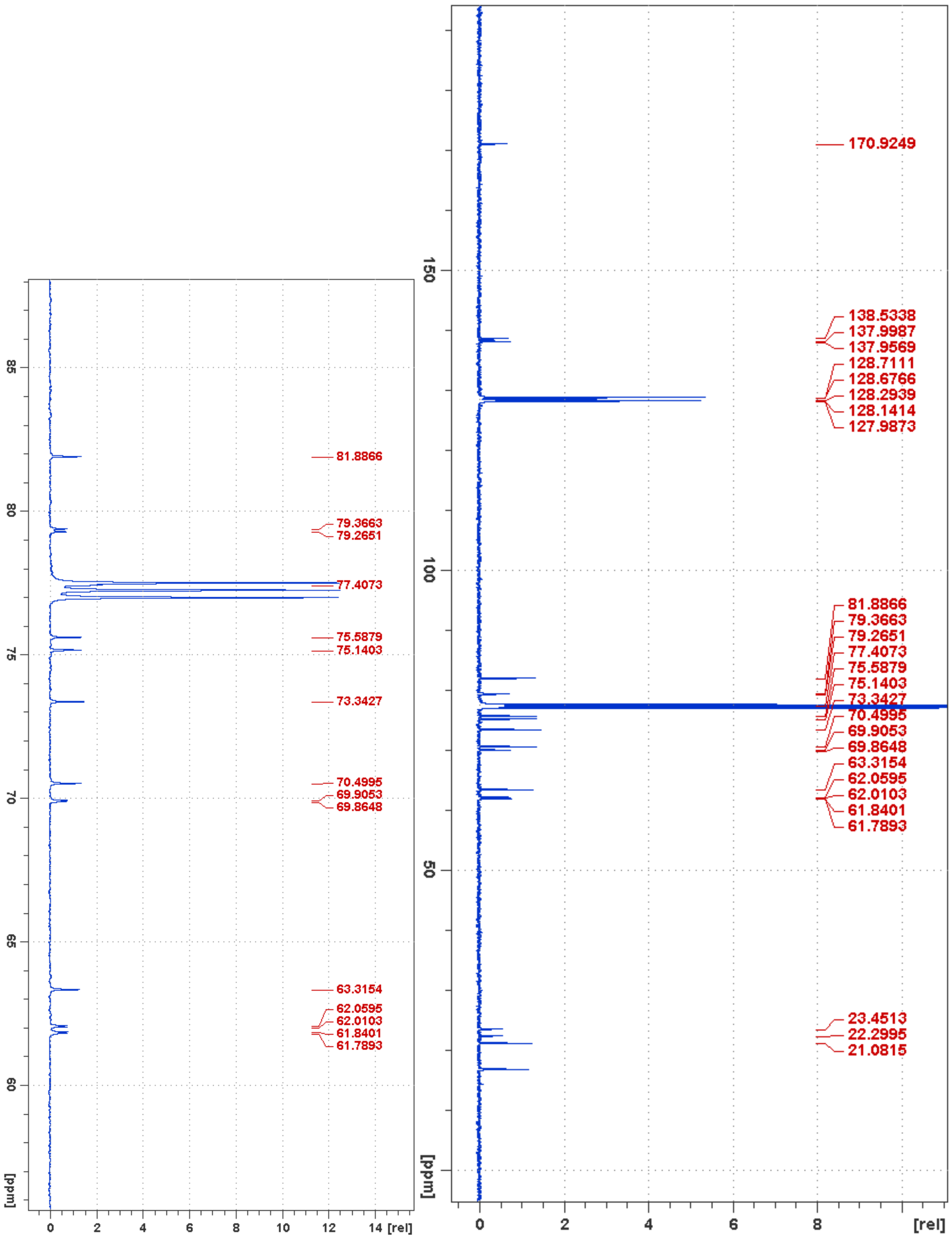


**Figure S12**. $^{13}$C NMR of compound **2**

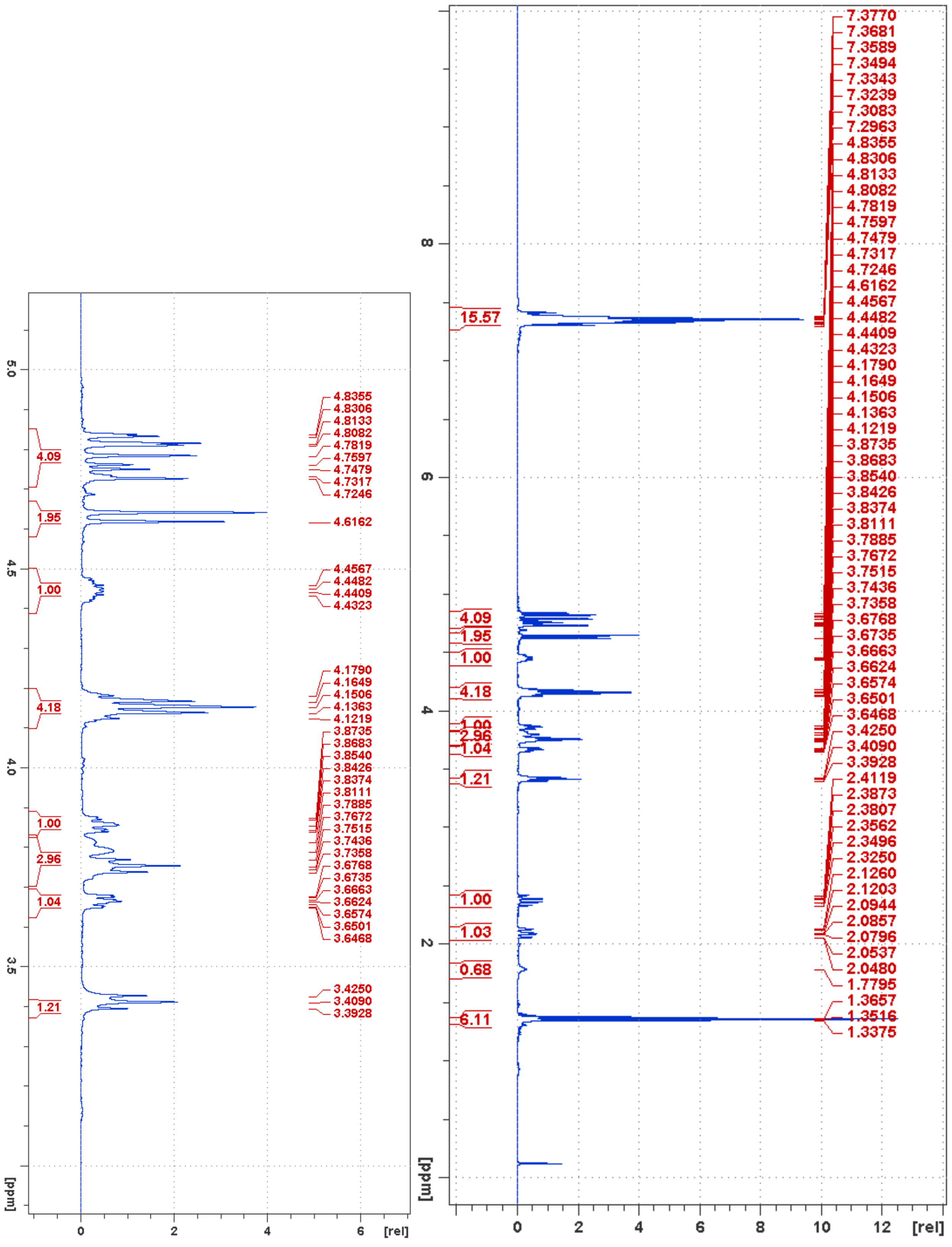


**Figure S13**. $^{1}$H NMR of compound **3**

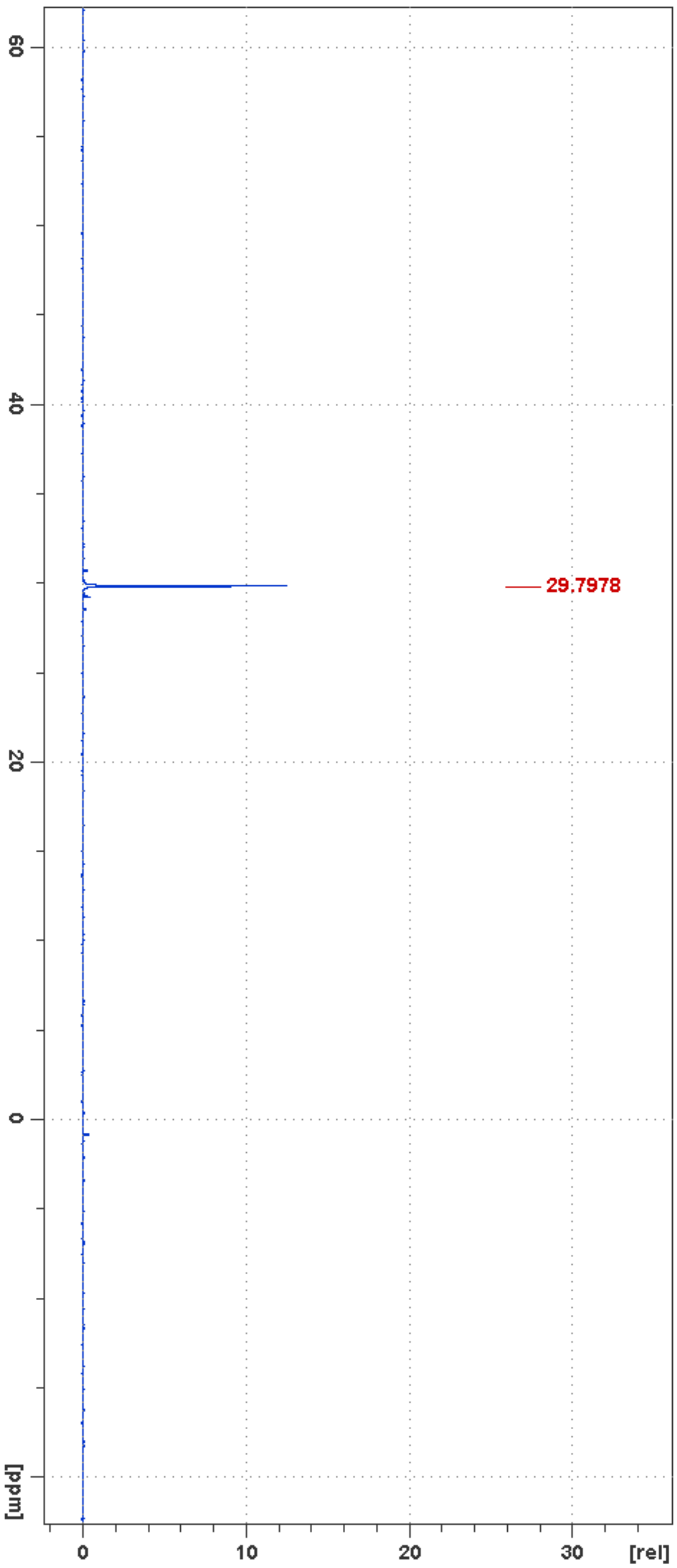


**Figure S14**. $^{31}$P NMR of compound **3**

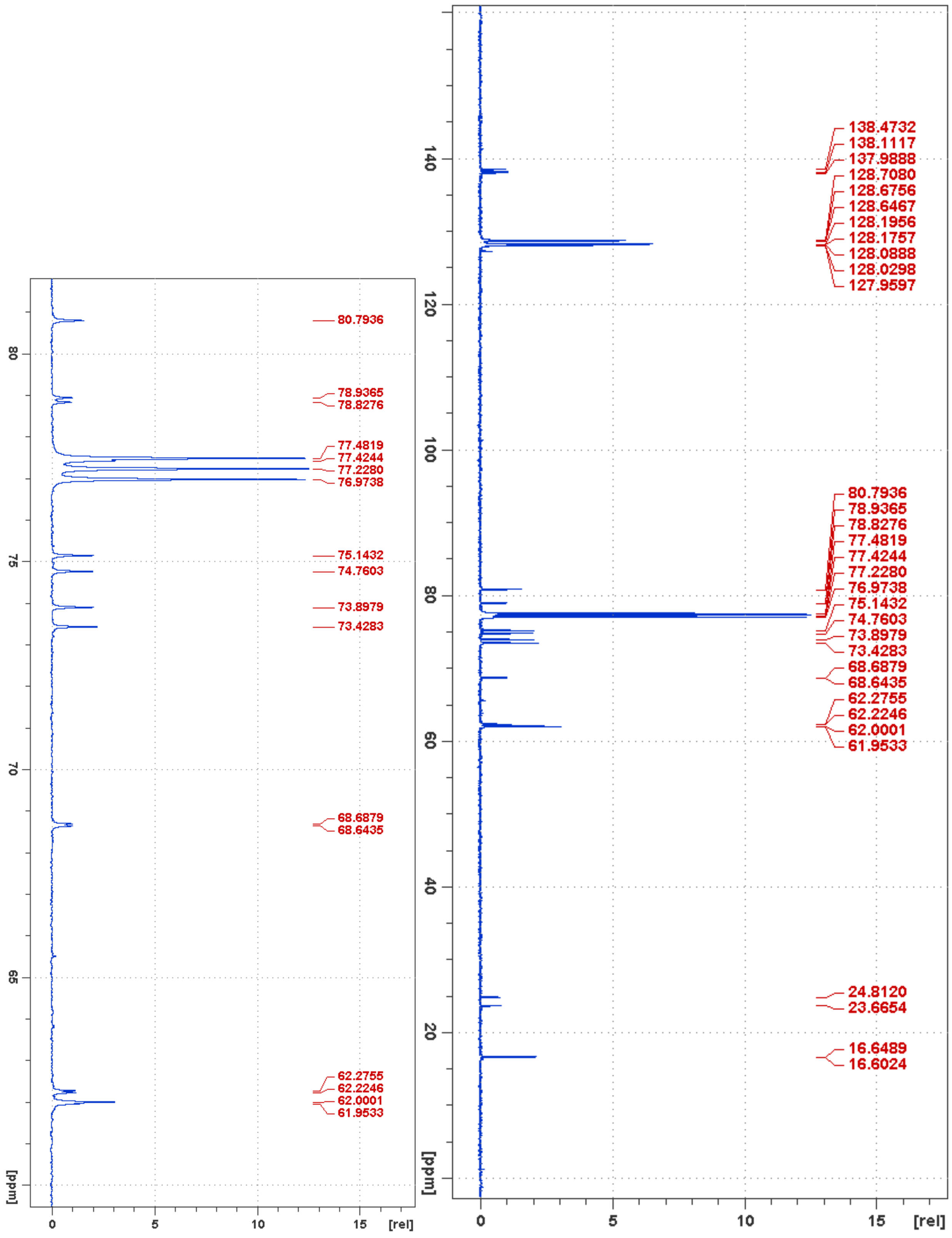


**Figure S15**. $^{13}C$ NMR of compound **3**

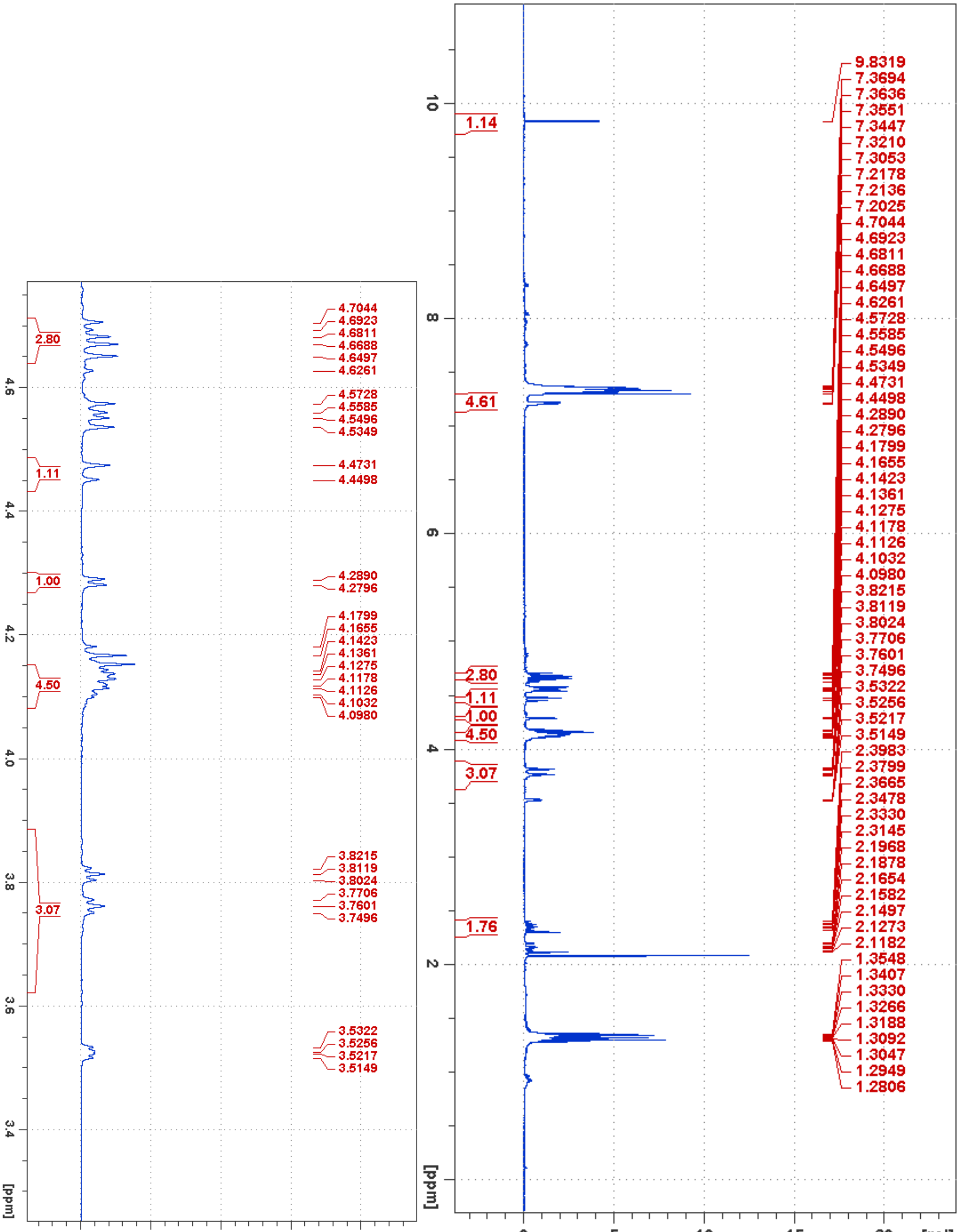


**Figure S16**. $^{1}H$ NMR of compound **4**

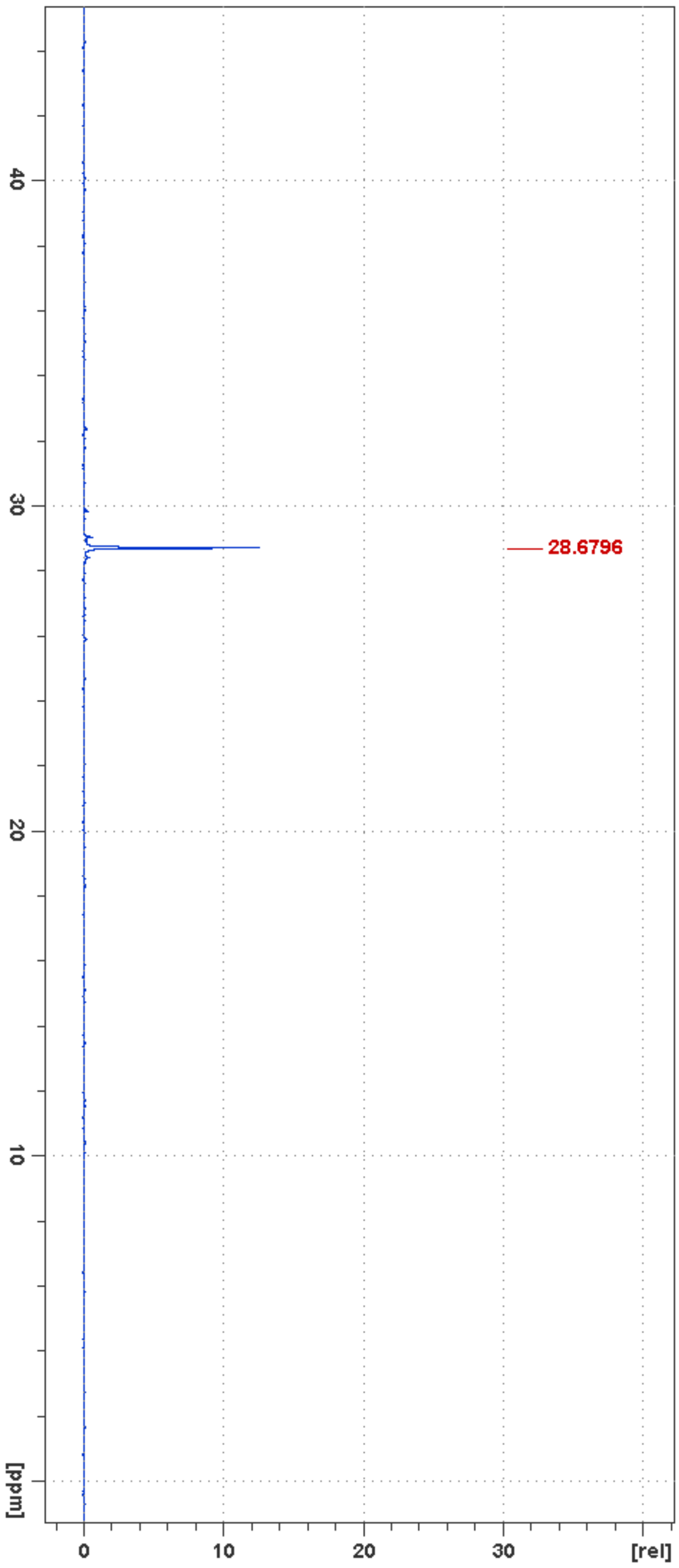


**Figure S17**. $^{31}$P NMR of compound **4**

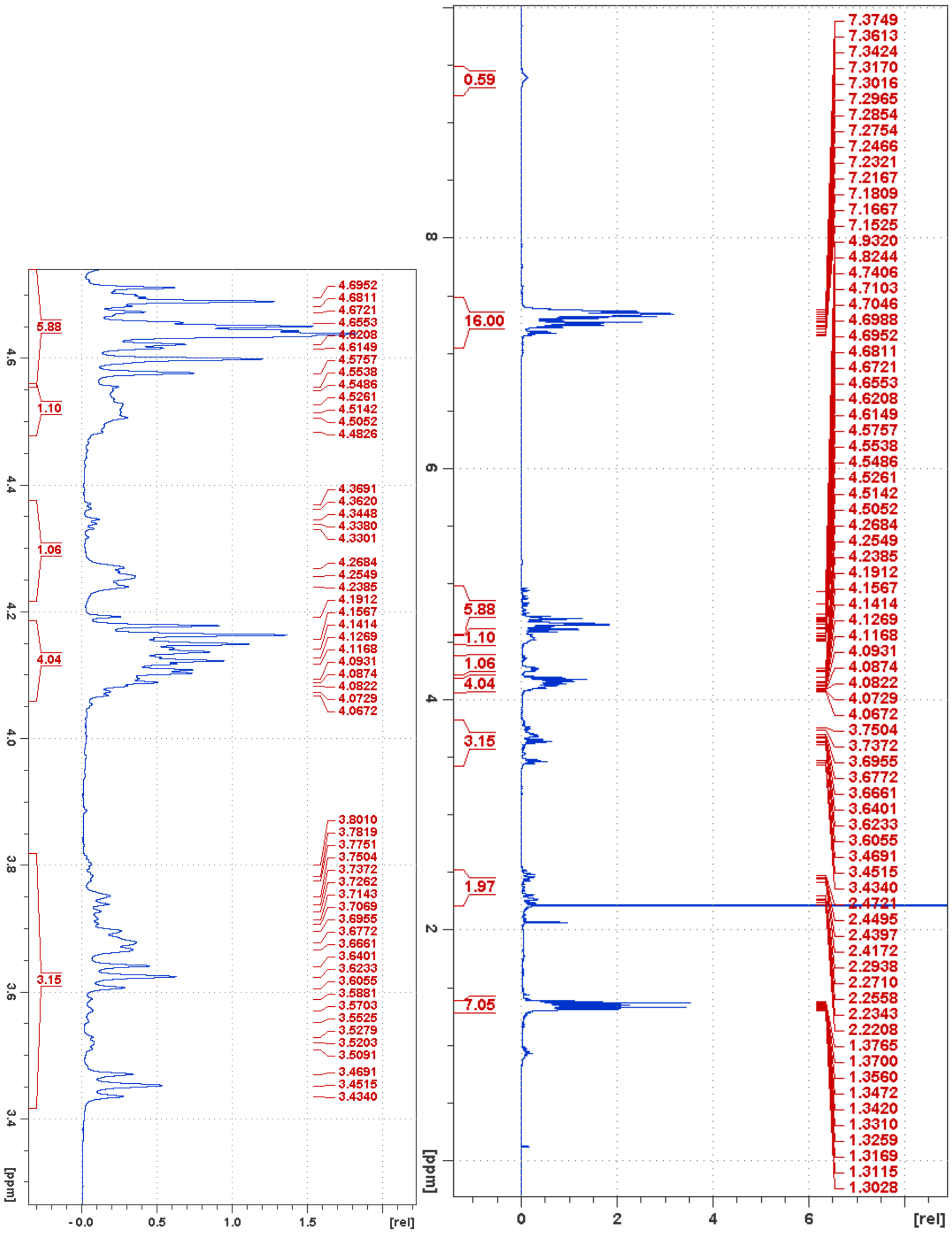

**Figure S18**. $^{1}$H NMR of compound **5**

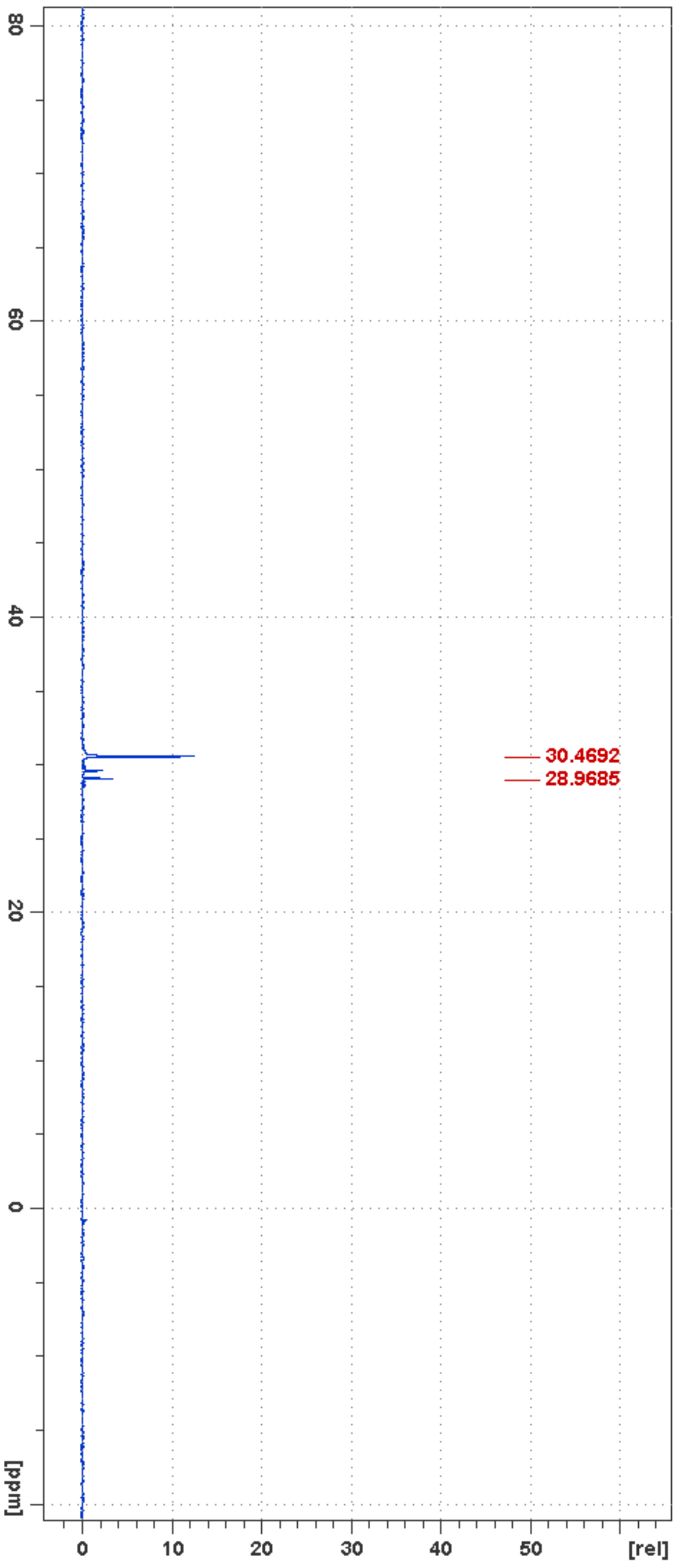


**Figure S19**. $^{31}$P NMR of compound **5**

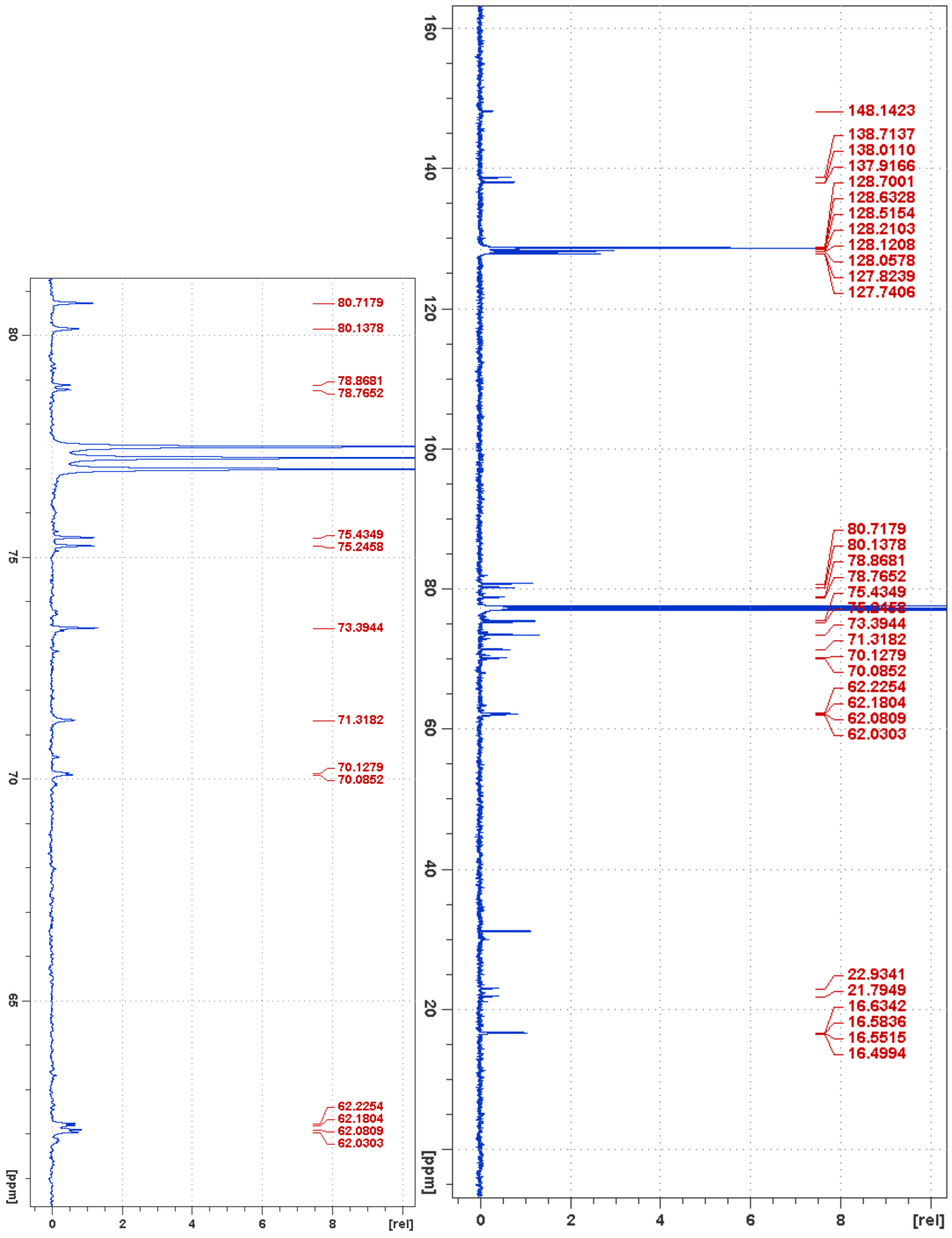


**Figure S20**. $^{13}$C NMR of compound **5**

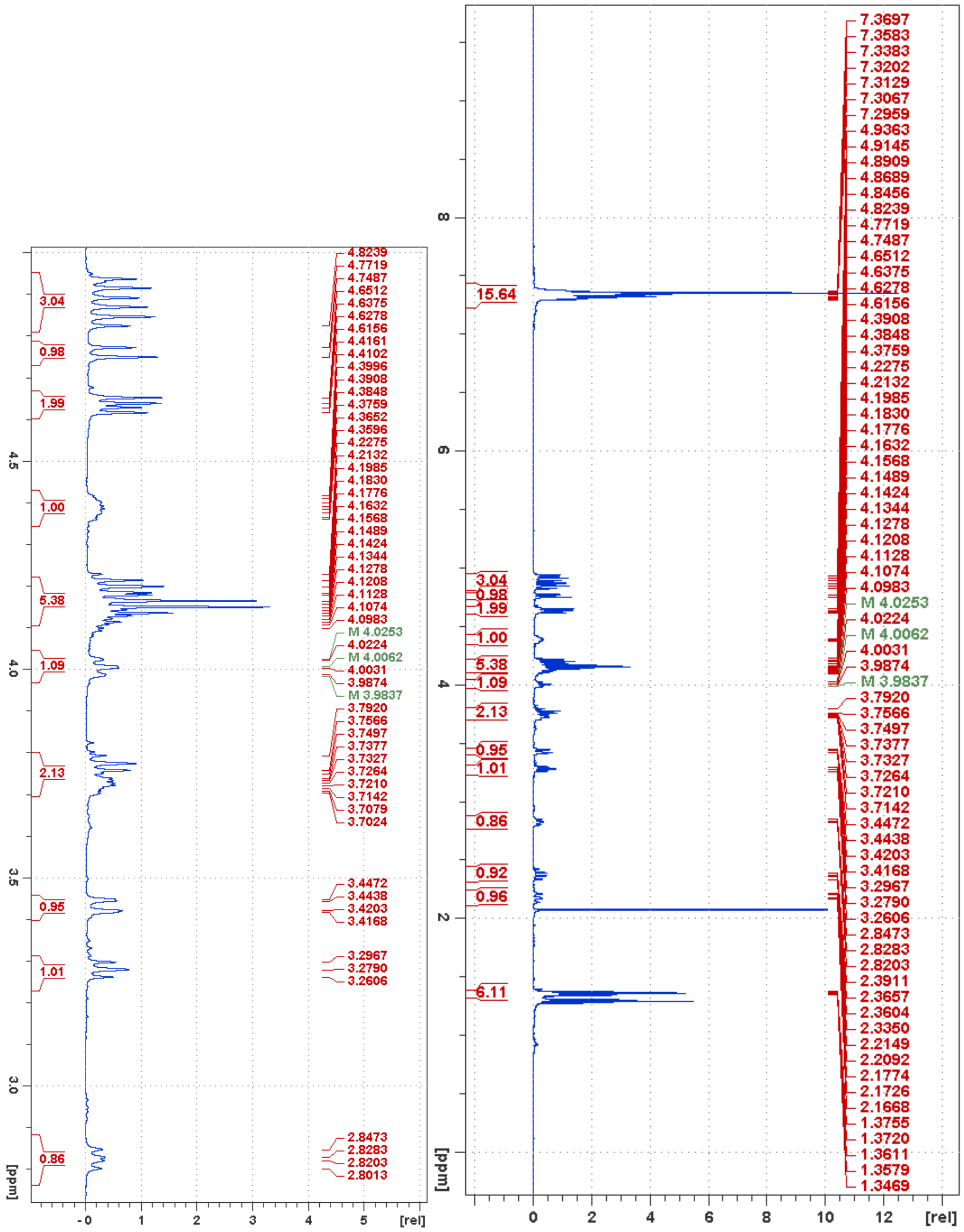


**Figure S21**. $^1H$ NMR of compound **6**

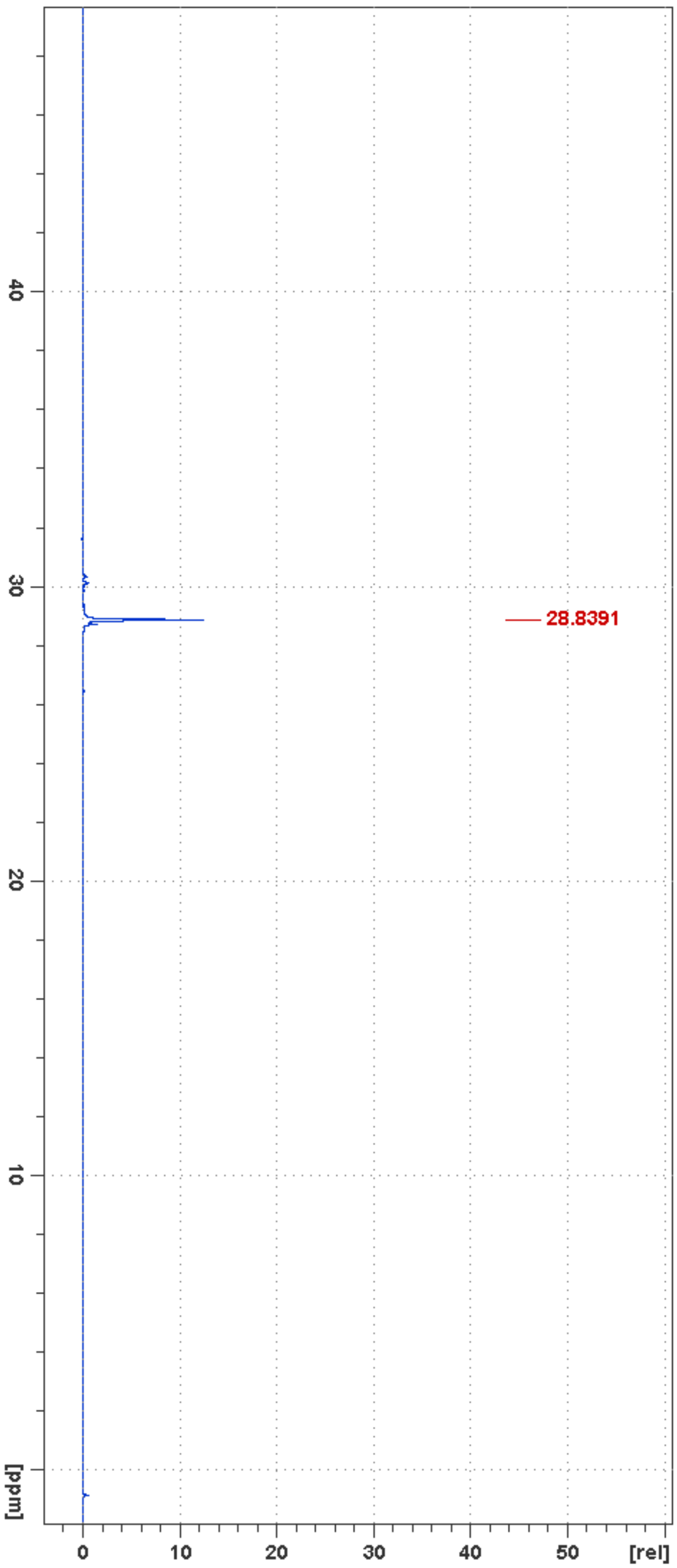


**Figure S22**. $^{31}P$ NMR of compound **6**

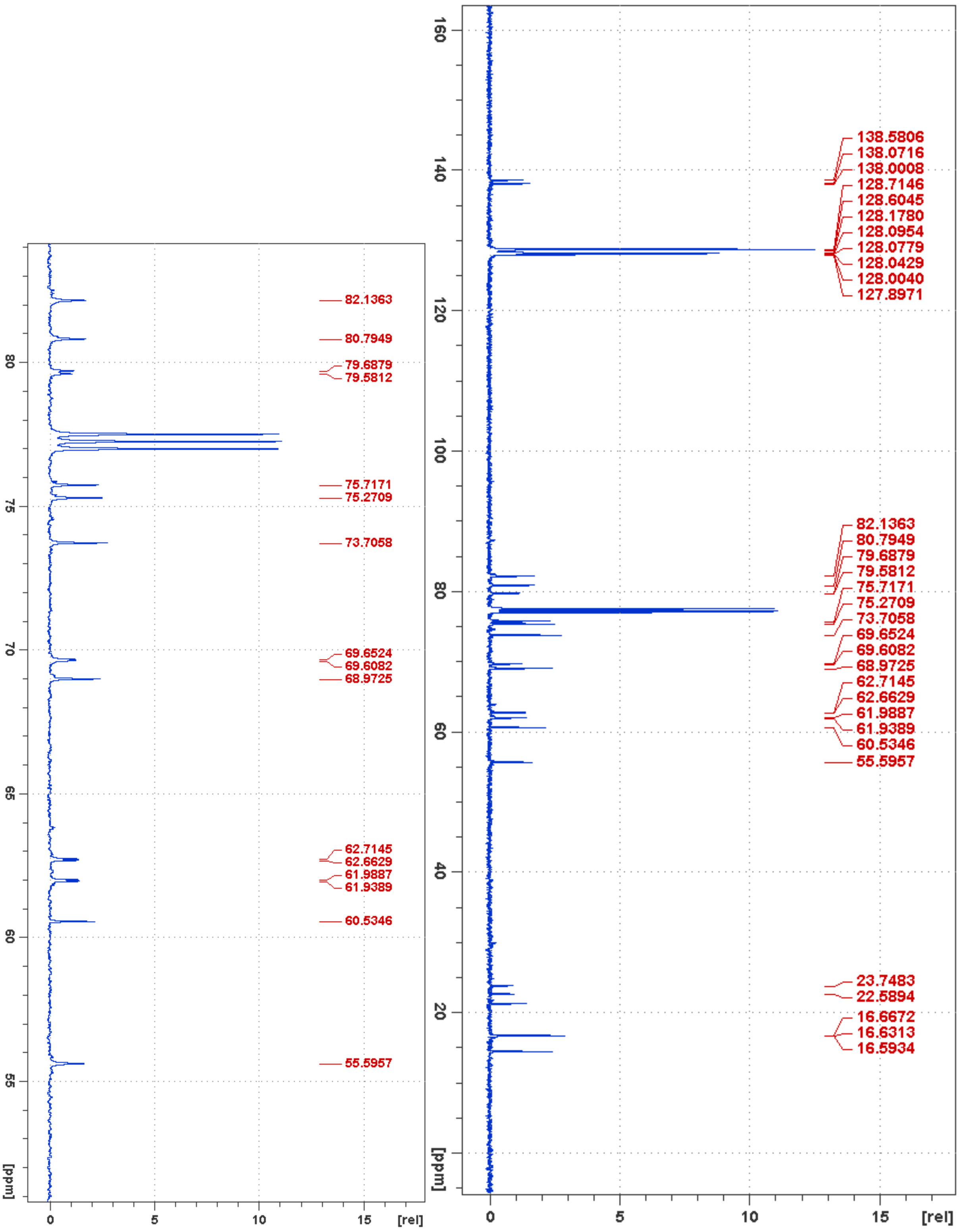


**Figure S23**. $^{13}$C NMR of compound **6**

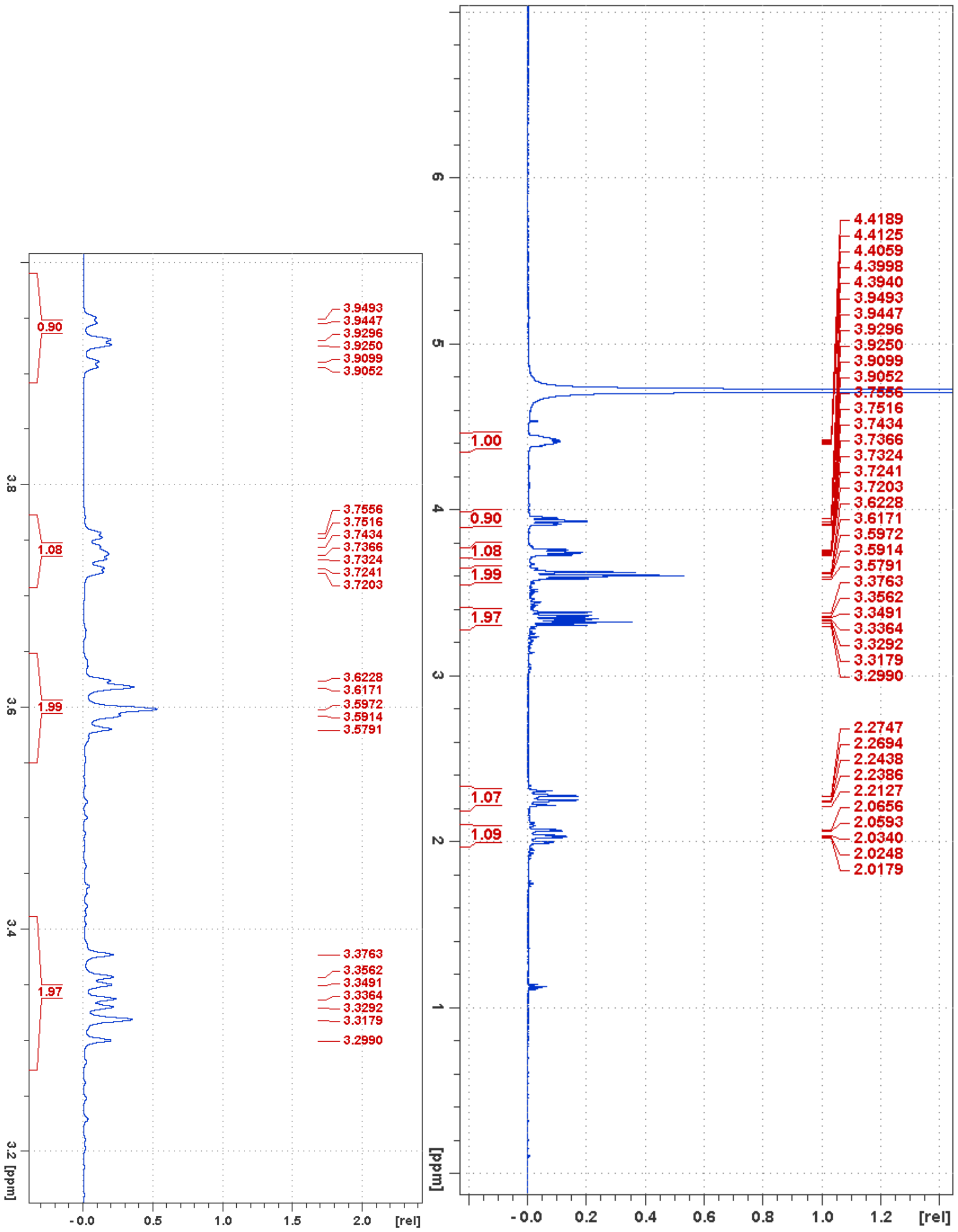


**Figure S24**. $^1$H NMR of compound **D9**

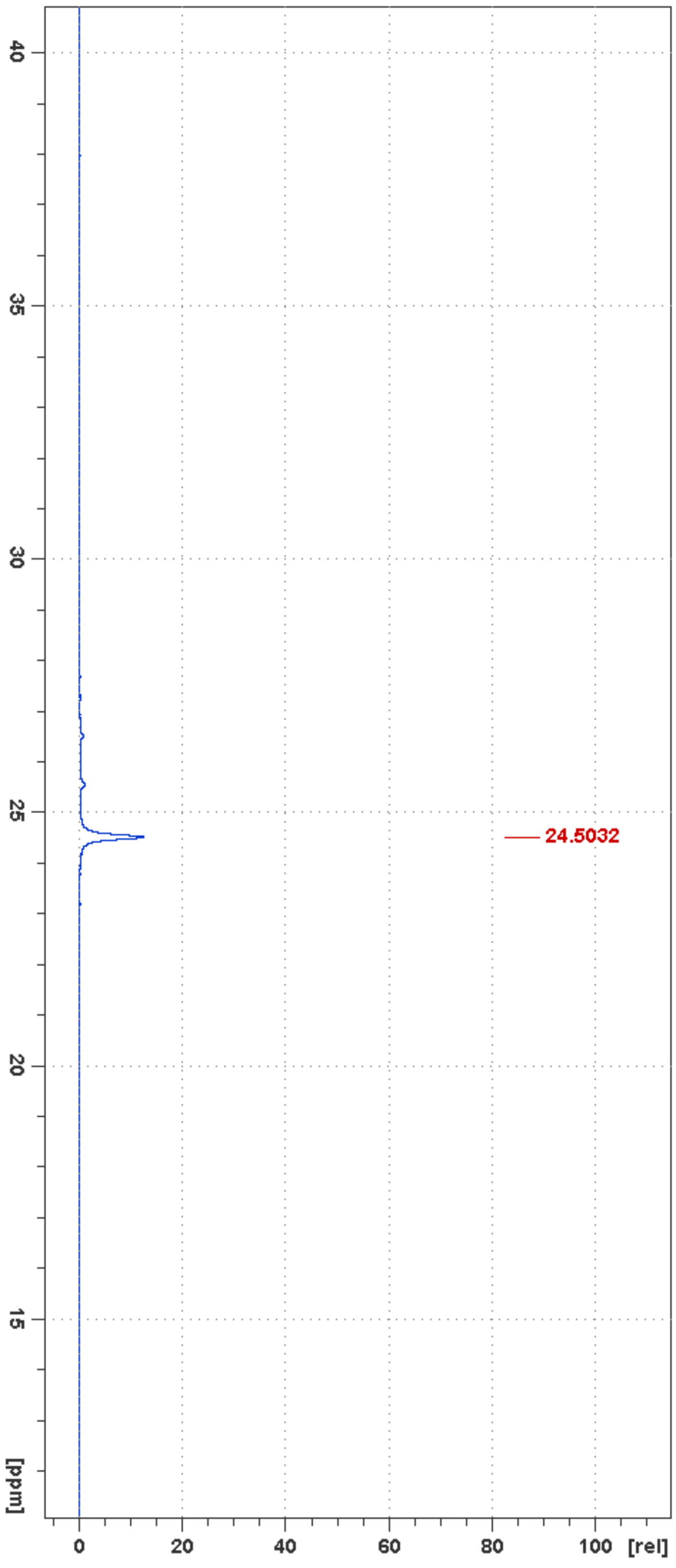


**Figure S25**. $^{31}P$ NMR of compound **D9**

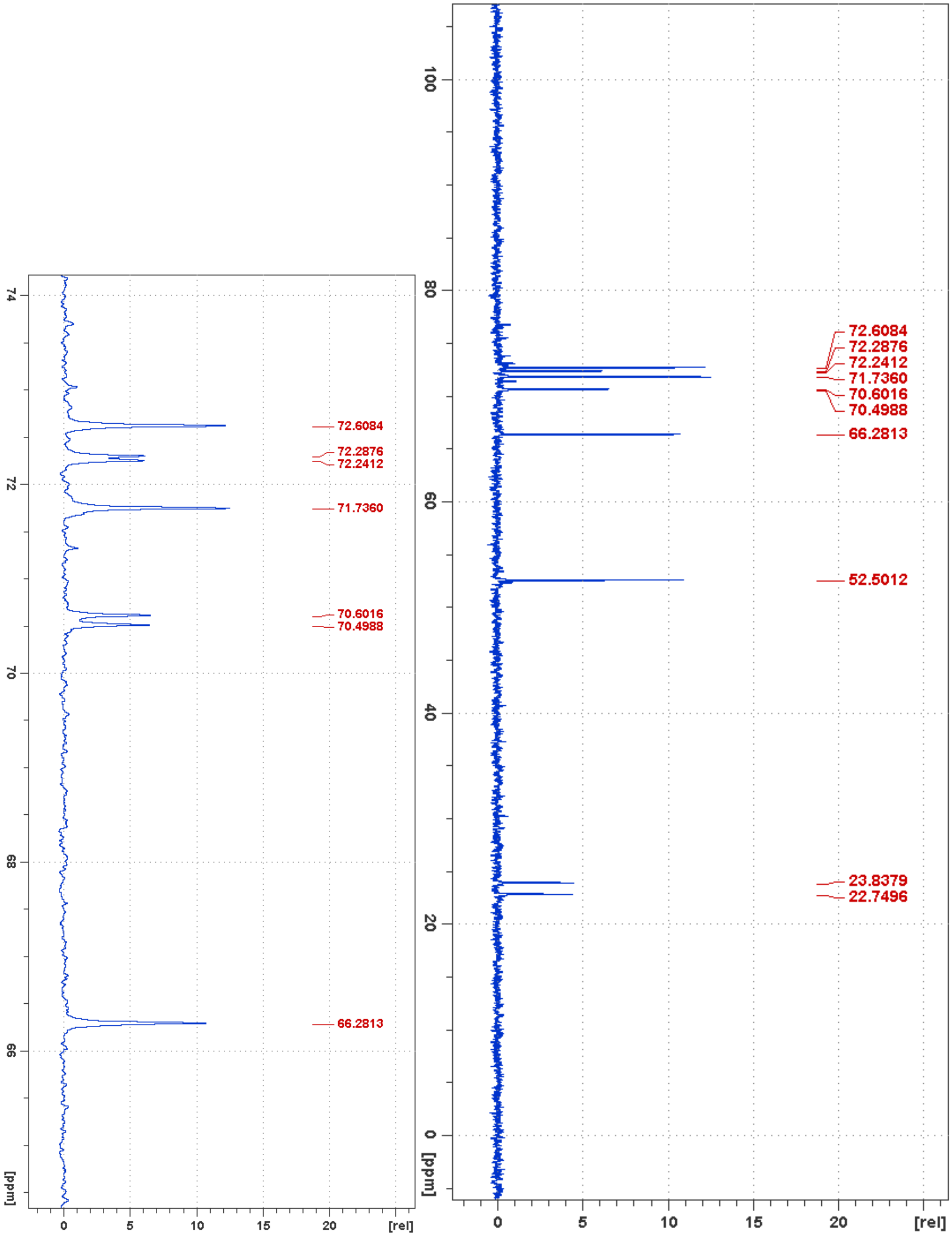


**Figure S26**. $^{13}C$ NMR of compound **D9**
